# Sleeping ribosomes: bacterial signaling triggers RaiA mediated persistence to aminoglycosides

**DOI:** 10.1101/2020.11.27.401281

**Authors:** Manon Lang, Evelyne Krin, Chloé Korlowski, Odile Sismeiro, Hugo Varet, Jean-Yves Coppée, Didier Mazel, Zeynep Baharoglu

## Abstract

Indole is a small molecule derived from tryptophan degradation and proposed to be involved in bacterial signaling. We find that indole secretion is induced by sublethal tobramycin concentrations and increases persistence to aminoglycosides in *V. cholerae*. Indole transcriptomics showed strongly increased expression of *raiA*, a ribosome associated factor. Deletion of *raiA* abolishes the appearance of indole dependent persisters to aminoglycosides, while its overexpression leads to 100-fold increase of persisters, and a reduction in lag phase, evocative of increased active 70S ribosome content, which was confirmed by sucrose gradient analysis. We propose that, under stress conditions, inactive 70S ribosomes are associated with RaiA to be stored and rapidly reactivated when growth conditions become favorable again, in a mechanism different than ribosome hibernation. Our results point to an active process of persistent cell formation, through ribosome protection during translational stress and relief upon antibiotic removal. Translation is a universal process, and these results could help elucidate a mechanism of persistence formation in a controlled, thus inducible way.

## Introduction

Antibiotic resistance is a major public health concern leading to increased health care costs and mortality (Opatowski et al., 2019; Touat et al., 2019). Although the majority of studies address the response of bacteria to lethal doses of antibiotics, the effect of low doses of antibiotics on bacteria has also recently started to draw attention. Antibiotic concentrations lower than the minimal inhibitory concentration (sub-MICs) have historically been proposed to serve as signaling molecules (Davies et al., 2006), provoking considerable changes in transcription and triggering a wide variety of cellular responses in different bacterial species (Andersson and Hughes, 2014) and mutagenesis (Gutierrez et al., 2013). In *V. cholerae*, sub-MIC aminoglycosides (AGs) are known to activate various stress response pathways, such as the SOS and RpoS stress responses, allowing to cope with increased reactive oxygen species (ROS) levels. AGs are bactericidal antibiotics that are known to enter the bacterial cell through the proton motive force (Fraimow et al., 1991; Herisse et al., 2017; Taber et al., 1987). AGs target the ribosome, leading to mistranslation and eventually cell death.

Interestingly, sub-MIC antibiotics, among which AGs, have been shown to stimulate the production of a small molecule, indole (Han et al., 2011). Indole is a byproduct of tryptophan degradation by tryptophanase TnaA (Evans et al., 1941) in more than 85 gram+ and gram - bacterial species (Lee and Lee, 2010), together with pyruvate and ammonia. While pyruvate and ammonia are respectively sources of carbon and nitrogen, the role of indole is not well understood. Indole is also found in plants, animals and was linked with signaling and human diseases (for a review, (Lee et al., 2015)). Common indole concentrations in the human gut are in the order of 250-1100 µM, and up to 200 µM in blood and other tissues. Regulation of indole production has been described, namely through carbon source utilization (Botsford and DeMoss, 1971) and catabolic repression (Yanofsky et al., 1991), aminoacid availability (Newton and Snell, 1965), cold temperature (Lee et al., 2008), heat shock (Li et al., 2003) and growth phase (Kobayashi et al., 2006). Particularly, indole is produced during transition from exponential to stationary phase (Lelong et al., 2007).

In *E. coli*, indole is nontoxic at physiologic concentrations (below 1 mM) (Lee et al., 2007), and does not change the growth rate (Lee et al., 2008). At high concentrations however (above 1-3 mM), indole inhibits cell division (Chant and Summers, 2007; Chimerel et al., 2012). In *V. cholerae*, indole secretion reaches its maximum at 600 µM during transition from midlog to stationary phase (Howard et al., 2019; Mueller et al., 2009) and was not observed to have any effect on the polarity of the *V. cholerae* cell membrane at this concentration (Mueller et al., 2009).

Indole can pass accross the cell membrane without the need for a transporter (Pinero-Fernandez et al., 2011), and was proposed to act as an interkingdom signaling molecule (Martino et al., 2003; Wang et al., 2001). An effect of indole in persistence to antibiotics has also been observed: it was found in *E. coli* that indole increases survival/persistence to lethal concentrations of ofloxacin, ampicillin and kanamycin (Lee et al., 2008; Vega et al., 2012), suggesting that the protective effect of indole is not specific to one family. Studies also pointed at the involvement of indole secretion in the cooperation between antibiotic resistant and sensitive populations during antibiotic stress (Lee et al., 2010). Notably, a recent study identified indole production as a potential target for the increased activity of quinolones against persisters in *E. coli* (Zarkan et al., 2020). Since indole appears to be beneficial for bacteria in presence of antibiotics, we addressed whether indole production is increased upon sub-MIC AG treatment in *V. cholerae* and whether this can lead to improved response to lethal antibiotic concentrations.

Importantly, we find that indole strongly increases persistence to AGs through the action of RaiA. We find that transcription from the *raiA* gene promoter is highly upregulated in the presence of indole in exponential phase *V. cholerae* cells. RaiA was shown to be a ribosome associated protein, in the same conditions as Rmf (ribosome modulation factor) and Hpf (hibernation promoting factor) (Maki et al., 2000). The two latter factors cause dimerization of vacant 70S into inactive 100S ribosome dimers, in a process called ribosome hibernation during stationary phase (Gohara and Yap, 2018), whereas RaiA mostly binds to free 70S monosomes (Maki et al., 2000; Sabharwal et al., 2015). *E. coli* mutants lacking these ribosome-associated factors do not show any growth defect during exponential growth, which is consistent with the fact that their expression is specific to stationary phase and stress (Prossliner et al., 2018). RaiA was observed to protect 70S ribosome from degradation (Agafonov et al., 1999; Di Pietro et al., 2013), and was also observed to block the binding of tRNA to the ribosomal A site in a cell free translation system (Agafonov et al., 2001), and during cold shock (Vila-Sanjurjo et al., 2004). In the present study, characterization of the *raiA* deletion mutant shows that RaiA is instrumental in the appearance of persister cells to AGs. We propose here a new mechanism of induced persistence to AGs by which RaiA positively affects the intact ribosome content of the cell, and facilitates regrowth after removal of the antibiotic.

## Results

### Indole is produced in response to sub-MIC tobramycin and increases persistence to AGs

Since the antibiotics ampicillin and kanamycin increase indole levels in *E. coli* (Han et al., 2011), we measured indole secretion (Saint-Ruf et al., 2014) in *V. cholerae* to determine whether the aminoglycoside tobramycin also impacts indole levels in this case. We found increased extracellular indole concentrations in the presence of sub-MIC tobramycin (TOB, from 20% of the MIC, **Figure S1A**). We next addressed whether indole has an impact on the growth of in *V. cholerae*, using an indole concentration of 350 µM. This concentration was previously shown to be physiologically relevant in *V. cholerae* and was observed to have no inhibitory effect on growth, and to complement the biofilm formation defect of a *tnaA* mutant deficient for indole production (Mueller et al., 2009). We found that 350 µM indole does not affect growth in the absence of antibiotics, but improves growth in sub-MIC antibiotics tobramycin, chloramphenicol and carbenicillin (**Figure S1BC** and not shown), which is consistent with previous observations.

We next addressed the effect of indole in the response to lethal concentrations of antibiotics, by measuring persister cells formation in *V. cholerae*. In order to do so, we adapted to *V. cholerae*, a protocol developed for *E. coli* (Ivan Matic and Wei-Lin Su, personal communication). Early exponential phase cultures were treated with lethal doses of antibiotics (5 to 10 times the MIC) for 20 hours. We first confirmed that CFUs that grow after 20 hours of antibiotic treatment are indeed persistent cells, by performing survival curves (**Figure S2A**). The profile of the curves we obtained showed a characteristic plateau after 6 hours, consistent with the formation of persister cells (Brauner et al., 2016). Furthermore, these cells were not resistant to the antibiotic. We thus carried on with the quantification of persisters at 20 hours of antibiotic treatment. We found that *V. cholerae* cultures grown in the presence of indole yielded higher numbers of persistent cells to AGs (**Figure 1AB**), as previously observed for *E. coli* treated with kanamycin and ofloxacin (Vega et al., 2012), but we observed no effect for persistence to carbenicillin (**Figure 1C**). Furthermore, a strain deleted for *tnaA* yielded less persisters to tobramycin than the WT strain (**Figure S2B**), and the effect was reversed by indole complementation, consistent with a link between indole and persistence.

**Figure 1.**
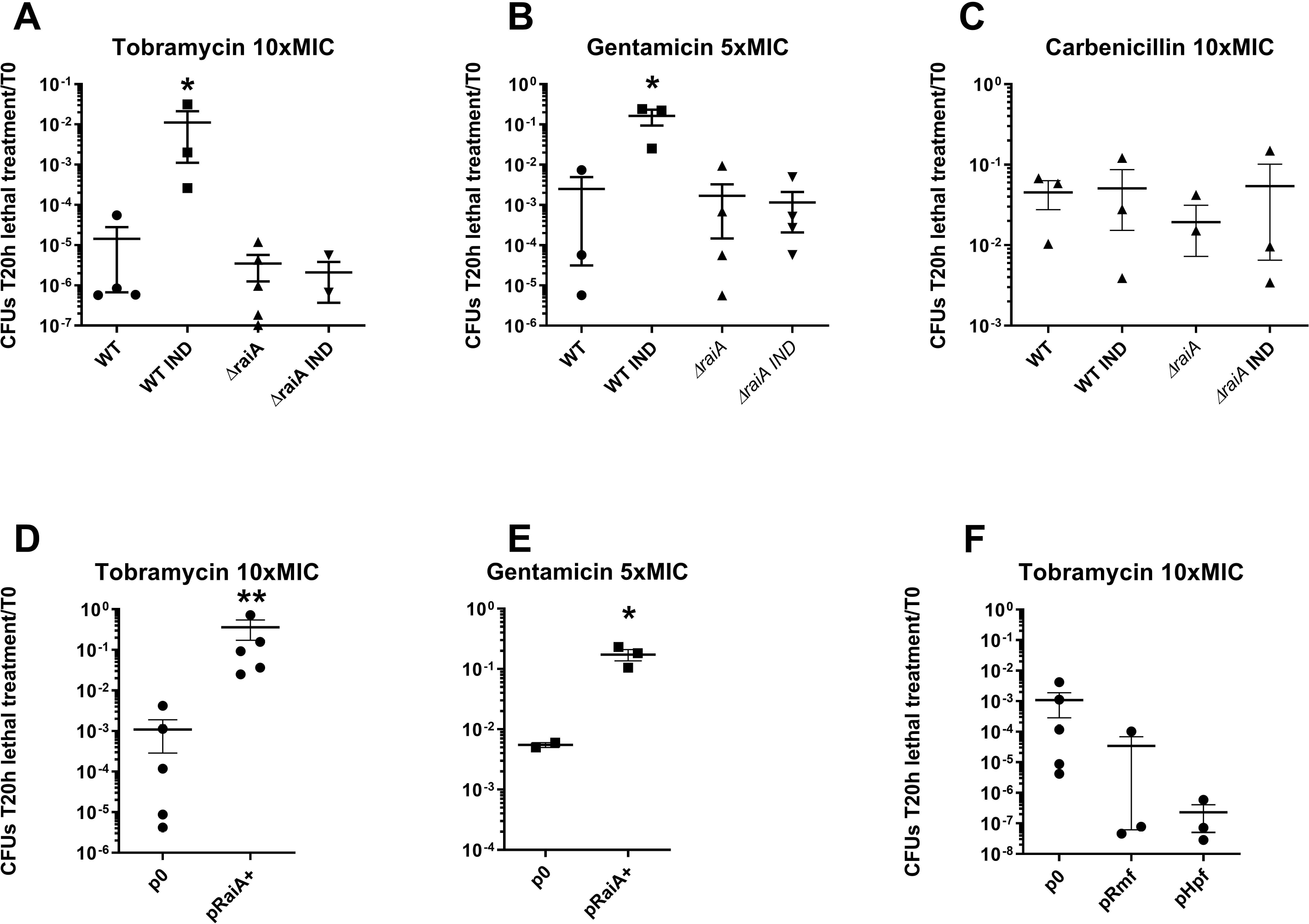
Modulation of persistence of exponential phase *V. cholerae* by indole and RaiA. Early exponential phase of wild-type (WT) and *ΔraiA V. cholerae* cultures were treated with lethal doses of the specified antibiotics for 20 hours, the ratio of CFUs growing after removal of the antibiotic and plating over total number of CFUs at time zero (before antibiotic treatment) are represented. Tobramycin (TOB): 10 µg/ml, gentamycin (GEN): 5 µg/ml, carbenicillin (Carb): 100 µg/ml, indole (IND): 350 µM. Experiments were performed 3 to 6 times, and statistical analysis was performed (t test, *: *p* <*0,05;* **: *p*<*0,005*). Experiments D, E and F were performed in WT strain carrying either the empty pBAD vector (p0) or with specified gene (pGene).

**Figure 2.**
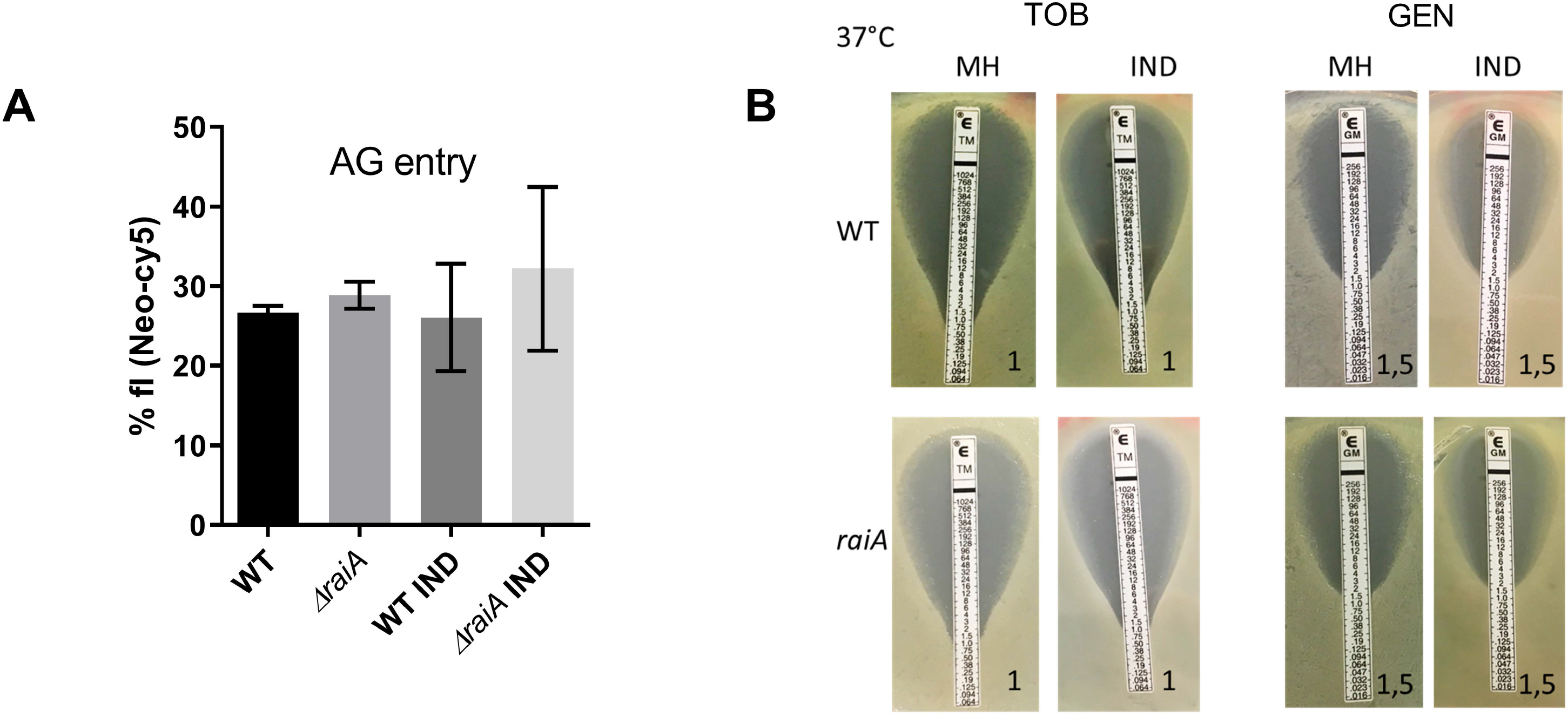
Indole does not influence aminoglycoside entry and resistance in *V. cholerae*. **A**. Intracellular level of neomycin coupled to the fluorophore Cy5 measured by fluorescence associated flow cytometry. **B**. Minimal inhibitory concentrations (MICs) of tobramycin (TOB) and gentamycin (GEN) measured using *etests* in *V. cholerae*, in the absence (MH) and presence of indole (IND), and indicated in µg/ml by a numeral on each image.

### Indole, at concentrations allowing growth, does not affect AG entry

AG uptake by the bacterial cell is known to be linked to the proton motive force (PMF). At high concentrations (5 mM) indole is a proton ionophore which blocks cell division by dissipating the PMF (Kralj et al., 2011), and was observed to interact with the cell membrane and change its physical structure (Mitchell, 2009). We thus addressed whether the beneficial effect of indole treatment in presence of AGs is due to modifications in membrane potential, which could lead to decreased AG entry into the cell. We used neomycin coupled to Cy-5 in order to measure AG entry in the bacterial cell (Sabeti Azad et al., 2020) (and Pierlé et al, unpublished), and found that indole at the physiological concentration of 350 µM does not affect AG entry into the bacterial cell (F**igure 2A**), ruling out the possibility of decreased AG entry due to modifications of PMF in presence of indole. Moreover, the presence of 350 µM indole did not change the MIC of two aminoglycosides tobramycin and gentamycin (F**igure 2B**). The beneficial effect of indole in presence of AGs is thus not through reduced antibiotic entry.

### Indole induces RaiA (VC0706) expression

In order to shed light into mechanisms allowing for more efficient response to antibiotic stress upon indole treatment, we decided to study the transcriptomic changes of *V. cholerae* in response to 350 µM indole. mRNA sequencing of exponential phase cultures shows differential regulation of 260 genes shown in **Table S1** (>2-fold change, adjusted p value <0.05 (Krin et al., 2018)). Notably, two translation related genes were markedly upregulated: *raiA* (20-fold up), together with *rmf* (3-fold up), which are both described as factors associated with inactive ribosomes in stationary phase (Agafonov et al., 1999; Di Pietro et al., 2013). *raiA* expression is known to be triggered by transition to stationary phase (Maki et al., 2000), and RaiA is usually weakly expressed during exponential phase. RT-qPCR on *raiA* and fluorescence associated flow cytometry (**Figure 3AB**) on cells carrying GFP fused to the *raiA* promoter confirmed upregulation of *raiA* by indole during exponential phase, and increased expression of *raiA* in stationary phase. Since transcriptomic data pointed RaiA as one of the most differentially regulated genes by indole, we next decided to address the contribution of RaiA in the indole associated phenotypes in *V. cholerae*.

**Figure 3.**
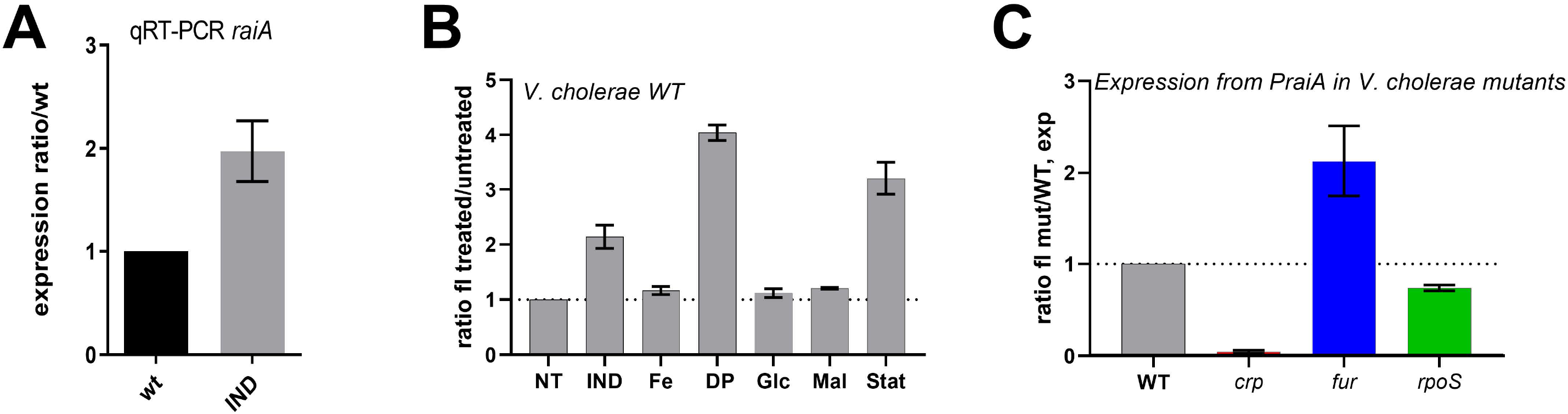
Expression of *raiA* in exponential phase *V. cholerae*. **A**. *raiA* mRNA levels measured by RT-qPCR on exponential phase *V. cholerae* cultures in presence or absence of indole. **B**. Fluorescence quantification of GFP expression from the *raiA* promoter by flow cytometry in MH media. NT: Non-treated, IND: indole (350 µM), Fe: iron (18 µM), DP: 2,2′-Dipyridyl (500 µM), Glc: Glucose (1%), Mal: Maltose (1%), Stat: stationary phase. **C**. Fluorescence quantification of GFP expression from the *raiA* promoter by flow cytometry in indicated *V. cholerae* deletion mutants. The Y axis represents fluorescence ratio of the mutant over wild type (WT) strain.

### Absence of *raiA* reduces persistence to AGs and prevents induction by indole

In order to address the involvement of RaiA in indole induced persistence, we constructed a *V. cholerae ΔraiA* mutant, and measured the appearance of persisters to tobramycin, gentamycin and carbenicillin in exponential phase cultures, grown in the absence and presence of indole (**Figure 1ABC**). We found that *V. cholerae ΔraiA* generally formed slightly less persistent cells to TOB, but the difference was not statistically significant, probably because at exponential phase *raiA* expression is not strong enough in the untreated WT strain. Strikingly, the deletion of *raiA* completely abolished induction of persistence by indole to both tested AGs (tobramycin, gentamycin, Figure 1AB), pointing to a role of RaiA in persister cell formation. No effect of neither indole nor *raiA* was observed in the formation of persisters to carbenicillin (**Figure 1C**), suggesting that the effect of RaiA in persister formation is specific to AGs. The involvement of RaiA in persistence to AGs also appears to be conserved in *E. coli* as the *raiA* deficient mutant yielded less persisters to tobramycin (here in stationary phase, **Figure S2C**). Finally, as performed above in the WT *V. cholerae* strain, we confirmed that the presence of indole does not affect the AGs MIC of the *ΔraiA* strain, and that deletion of *raiA* does not affect the MIC (**Figure 2**), meaning that the phenotypes we observe are not due to reduced entry of AGs or increased resistance. On the other hand, RaiA is dispensable for growth improvement by indole (**Figure** S**1D**), because indole still improves growth in TOB when *raiA* is deleted, showing that the mechanism of AG persister induction by indole (antibiotic concentration >MIC) is different than the mechanism of growth improvement in sub-MIC AGs. It is worth mentioning here that the growth of *ΔraiA* strain appeared to be slightly slower than the WT strain, suggesting that RaiA may also have a role during exponential growth.

### RaiA overexpression increases persistence to AGs and promotes earlier exit from stationary phase

In order to overexpress RaiA, we cloned it under a controlled pBAD promoter, which is repressed by glucose and induced by arabinose. Since the presence of different carbon sources may differentially affect growth and the response to aminoglycosides (Pierlé et al., unpublished), we compared the persistence levels of cells carrying the empty vector (p0 in Figure 1DE) or the pBAD-RaiA plasmid, in the presence of arabinose (RaiA overexpression conditions). Overexpression of RaiA strongly increased persister formation in tobramycin and gentamycin (**Figure 1DE**), indicating that RaiA is directly involved in the persistence mechanism.

We next asked whether such increased persistence could be due to slower growth when RaiA is overexpressed. We monitored growth in conditions where 1) RaiA is not overexpressed previous to inoculation and only overexpressed during growth, and 2) RaiA is previously overexpressed in cells used for inoculation. First, our results show no difference in growth rate (slope) in presence or absence of RaiA overexpression (F**igure 4AB**), indicating that RaiA overexpression does not cause a slow growth phenotype, which discards the hypothesis linking RaiA-mediated persistence to slow growth/dormancy. Interestingly, we observe in cells were RaiA overexpression was pre-induced, that these cells start growing faster due to a reduction in lag phase (**Figure 4CD**). Such a reduction in lag phase is reminiscent of increased active ribosome content which allows faster resumption of growth at the exit of stationary phase (Condon et al., 1995). As a corollary, when *raiA* was deleted, the lag phase was increased (**Figure 4E**), suggesting that RaiA levels affect inactive but “ready to use” ribosome content in stationary phase.

**Figure 4.**
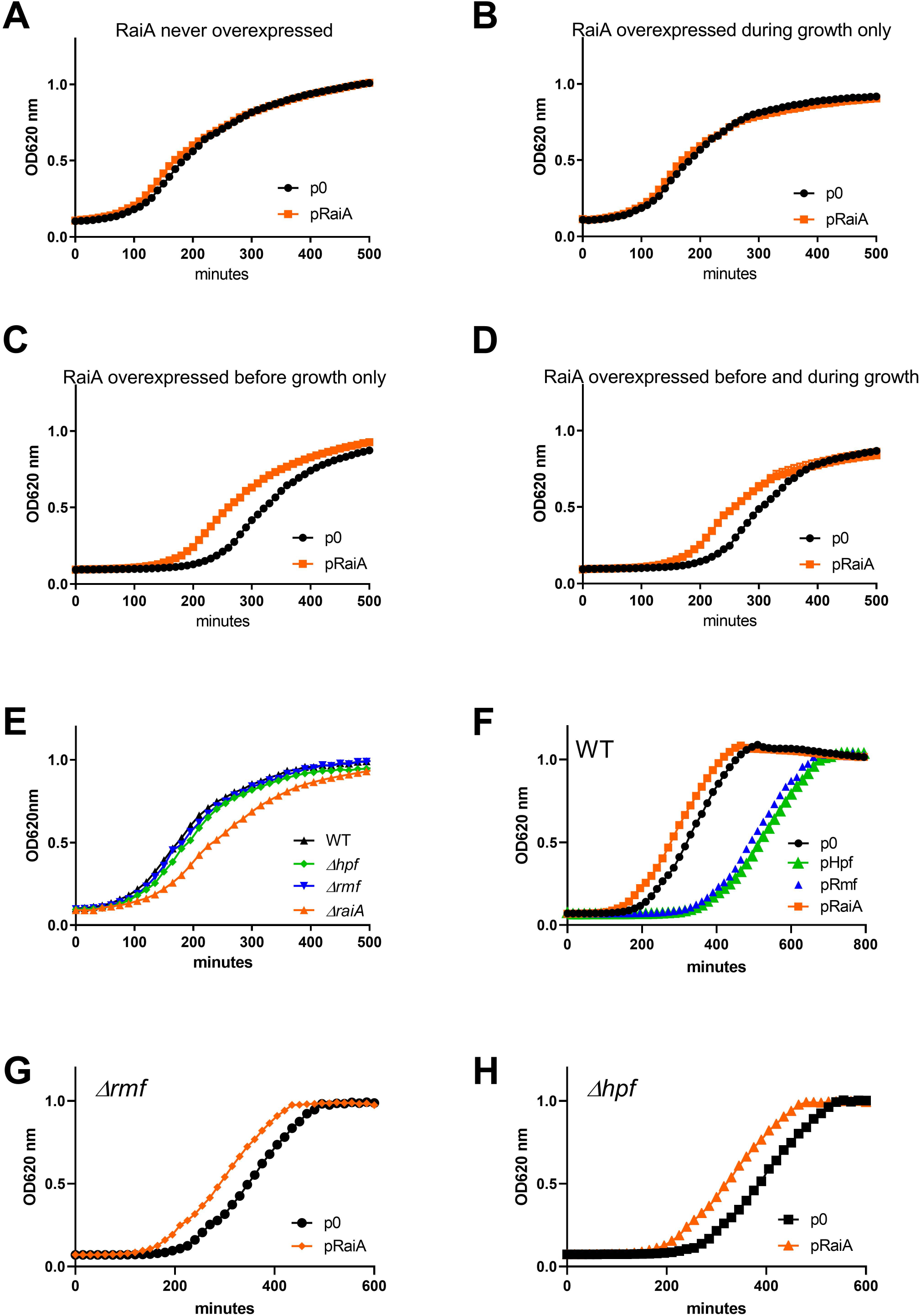
RaiA influences lag phase upon growth restart after the stationary phase. Growth is measured on a plate reader by measuring OD at 620nm (y axis), for the time indicated by the x axis. **ABCD**: The curves of wild type *V. cholerae* with the empty plasmid is compared with plasmid carrying *raiA* under inducible promoter. The promoter is repressed using glucose (GLC) and induced using arabinose (ARA). Growth was performed as specified, with RaiA expression repressed or induced in the overnight culture used for inoculum, and during growth in the plate reader, as specified (inoculum/growth): **A**. GLC/GLC, **B**. GLC/ARA, **C**. ARA/GLC, **D**. ARA/ARA. **E**. Growth in MH of WT and mutant strains. **F**. Growth of *V. cholerae* overexpressing hibernation factors or empty plasmid (p0). **GH**: Growth of hibernation factor mutants Δ*rmf* and Δ*hpf* overexpressing RaiA.

### The effect of RaiA on lag phase and persistence is independent of ribosome hibernation factors Rmf/Hpf

RaiA was previously found to be associated with the inactive ribosomes at stationary phase, in the same conditions as Rmf and Hpf factors. Rmf and Hpf are known to dimerize ribosomes into so called hibernating 100S ribosomes, whereas RaiA associates with monomeric 70S ribosomes (Gohara and Yap, 2018; Maki et al., 2000; Prossliner et al., 2018). In order to address whether Hpf/Rmf dependent ribosome hibernation also favors rapid exit from stationary phase, we performed deletion and overexpression experiments similar to what we did for RaiA. Deletion of *rmf* or *hpf* has no effect on lag phase nor growth (**Figure 4E**). Our results showed that in contrast to overexpression of RaiA which decreases the lag phase, overexpression of Rmf or Hpf rather increase lag phase (F**igure 4F**). These findings are consistent with a model where increased ribosome dimerization by Hpf/Rmf would require action of dissociation factors in order to resume growth whereas spontaneous dissociation of RaiA from inactive 70S ribosomes is sufficient for growth restart. Furthermore, when we overexpressed RaiA in Δ*hpf* or Δ*rmf* mutants, we observed a reduction of lag phase similar to what is observed upon overexpression of RaiA in the WT strain (**Figure 4GH**), meaning that RaiA action is not dependent of Hpf/Rmf mediated ribosome hibernation.

In addition, we addressed the effect of hibernation factors on persistence. Unlike for RaiA overexpression, no increase in persistence to TOB was detected upon overexpression of Hpf or Rmf, excluding an effect of 100S ribosome dimer formation on persistence to AGs (F**igure 1F**). Unexpectedly, persistence levels even decreased upon overexpression on the hibernation factors, suggesting that 70S-RaiA and 100S-Rmf/Hpf complexes have opposite effects on persistence to AGs. We also tested the persistence levels of *rmf* and *hpf* deletion mutants. Surprisingly again, and consistent with overexpression results, we found that deletion of *rmf* or *hpf* hibernation factors increases persistence to AGs (F**igure S2D**). The increase of persistence due to deletion of *rmf* is dependent on the presence of *raiA*, as the double mutant *raiA rmf* shows reduced persistence compared to WT, and similar to the *raiA* mutant. This can be due to the observed increased expression of *raiA* in the absence of *rmf* (Sabharwal et al., 2015) or amplified ribosome-RaiA complex formation in the absence of *rmf*, due to decreased 100S formation (Ueta et al., 2005). Finally, it is important to note that the *hpf raiA* double mutant could not be constructed despite the use of two different strategies, implying synthetic lethality. Overall these results suggest an equilibrium between Rmf/Hpf and RaiA actions, consistent with previous literature that showed a combined role for these proteins in ribosome hibernation and antagonizing regulation of *rmf/hpf* and RaiA in *V. cholerae* (Sabharwal et al., 2015).

### RaiA overexpression increases 70S ribosome proportion over 50S and 30S subunits in stationary phase

In order to address whether intact 70S ribosomes are protected/stored upon RaiA overexpression, we measured the ribosome contents of WT and RaiA overexpressing cells, as well as the strain deleted for *raiA*, by performing 10-50% sucrose gradients on cellular extracts from 24h stationary phase cultures. The profiles obtained were consistent with well described peaks for 30S and 50S subunits followed by a third peak corresponding to the 70S ribosome (Ueta et al., 2013). Deletion of *raiA* leads to a slight increase in the proportion of dissociated subunits compared to 70S ribosome. Such minor effect was probably due to low expression of RaiA in the WT strain in exponential phase. We found that the proportion of 70S ribosomes is increased compared to 30S+50S dissociated subunits upon RaiA overexpression (**Figure 5**). We observed similar profiles when the cultures were treated with lethal concentrations of TOB (**Figure S3**). This agrees with the hypothesis that RaiA would stabilize the intact 70S ribosome and increase functional ribosome pools.

**Figure 5.**
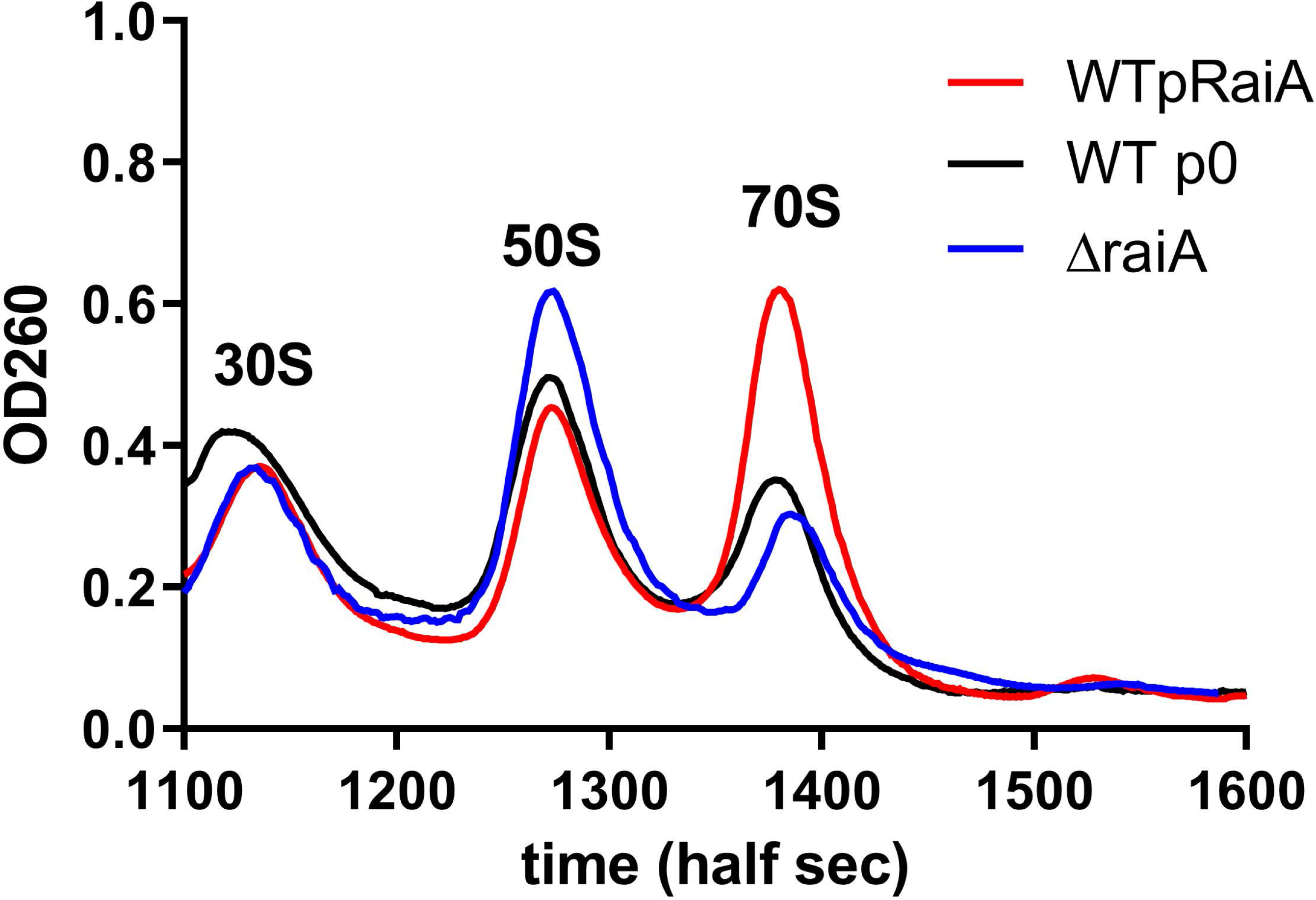
RaiA levels influence stationary phase 70S ribosome content relative to 50S+30S subunits in *V. cholerae*. Cellular extracts of 24 hours cultures were separated on 10-50% sucrose density gradient. Ribosomal RNA content was measured at OD 260nm using a spectrometer coupled to a pump and time on X axis represents samples from less dense (upper fragments, smaller complexes) to denser (bottom of the tube, heavier complexes). Cell debris eluting before 1100 half seconds are not shown. Graphs are normalized to total OD260nm=1 for each sample.

### RaiA expression is under environmental control and linked to the Fur iron sensing regulon in *V. cholerae*

RaiA is known to be expressed in stationary phase and upon temperature stress (Agafonov et al., 1999; Agafonov et al., 2001; Maki et al., 2000; Sabharwal et al., 2015; Slamti et al., 2007). Regulation of *raiA* was also described to occur through carbon catabolite control (CRP-cAMP) (Manneh-Roussel et al., 2018; Shimada et al., 2013). Our transcriptomic data suggests that indole affects genes from the Fur regulon (**Table S2**). Fur responds to iron levels and is generally known to be a repressor of iron/heme transport genes (such as *hutA, hutXW, tonB, fbpA, viuB*, among others) and also activates a small number of genes (namely the *napABC* operon and *menB* in *V. cholerae*) (Mey et al., 2005). The *V. cholerae raiA* gene promoter, appears to carry sequences similar to Fur boxes described in *V. cholerae* (Davies et al., 2011). In order to shed light in the means by which indole induces expression from the *raiA* promoter in exponential phase, we constructed deletion mutants for *fur, crp* and for the stationary phase sigma factor *rpoS*. We used our *PraiA-gfp* transcriptional fusion to measure expression in the presence and absence of indole in the mutants compared to wild type strain.

As expected, in WT *V. cholerae*, fluorescence was triggered by indole at 3 hours of culture which corresponds to early exponential phase (OD 0,2 to 0,3), and accumulated in stationary phase (**Figure 3B**). Deletion of *crp* strongly decreases *raiA* expression (25x in exponential phase, **Figure 3C**), confirming that CRP is a prevailing activator of *V. cholerae raiA*. On the other hand, deletion of *rpoS* had no major effect on *V. cholerae raiA* expression (1,25x decrease). In the *Δfur* strain, *raiA* expression was increased 2fold (Figure 3C), suggesting Fur dependent repression of the *raiA* promoter. No major effect on *raiA* promoter was observed upon treatment with iron, possibly because iron is already in excess levels for the cells during growth. Strikingly, treatment with dipyridyl (DP), an iron chelator which mimics conditions of iron starvation strongly induced fluorescence (**Figure 3B**).

These results show a link between extracellular iron levels and RaiA expression. Together with the CRP-cAMP control, RaiA expression appears to be under environmental control, highlighting a link between bacterial persistence and environmental stress.

### The effect of RaiA overexpression is conserved among Gram-negative pathogens

We next asked whether RaiA could have a similar function in other Gram-negative pathogens such as *Pseudomonas aeruginosa*, an organism associated with antibiotic resistance and persistence (Koeva et al., 2017; Spoering and Lewis, 2001) and of high concern regarding resistant infections. *P. aeruginosa* RaiA exhibits respectively 37% and 31% protein identity with RaiA from *V. cholerae* and *E. coli*. We overexpressed *V. cholerae* RaiA in *P. aeruginosa* and assessed lag phase and persistence, as we performed for *V. cholerae* (**Figure 6**). We found that upon RaiA overexpression, lag phase is also decreased in *P. aeruginosa*, and strikingly, persistence to tobramycin is increased. These results show that RaiA mediated ribosome protection can be involved in persistent infections by various pathogenic bacteria.

**Figure 6.**
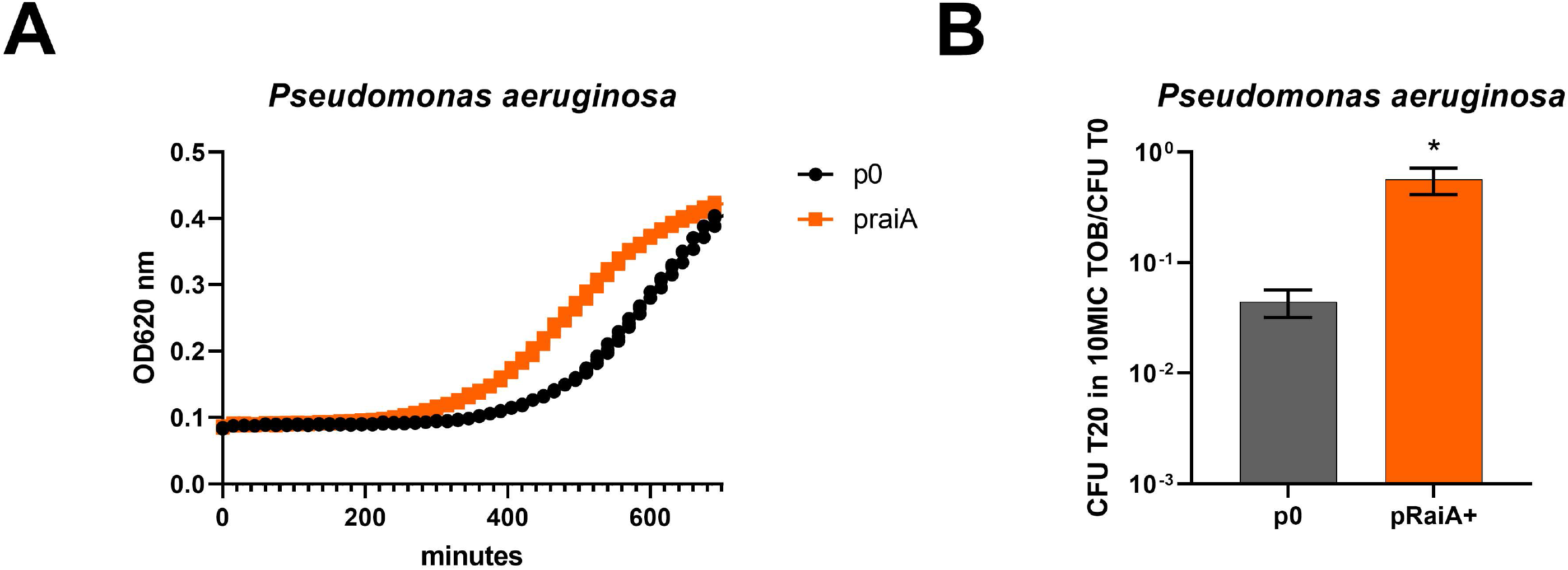
RaiA overexpression in Pseudomonas aeruginosa increases tolerance to tobramycin and promotes earlier exit from stationary phase. **A**. Growth is measured on a plate reader by measuring OD at 620nm (y axis), for the time indicated on the x axis. B. *Pseudomonas aeruginosa* carrying the empty pBAD vector (p0) or the pBAD vector overexpressing *raiA* of *V. cholerae* (*pRaiA+*), cultures were treated with lethal doses of tobramycin (10 µg/ml) for 20 hours, the ratio of CFUs growing after removal of the antibiotic and plating over total number of CFUs at time zero (before antibiotic treatment) are represented. (t test, *: *p* <*0,05*).

## Discussion

We show here that indole is produced upon sub-MIC aminoglycoside treatment in *V. cholerae* and at physiological concentrations, increases appearance of persister cells in lethal concentrations of AGs. We find that such increase in persistence occurs through an inducible mechanism involving RaiA (previously called pY or *yfiA* in *E. coli*, and *vrp* in *V. cholerae*). While indole improved growth seems to be nonspecific to an antibiotic class and independent of RaiA, increased persistence involving RaiA is specific to AGs. Since RaiA is regulated by several environmental cues and signaling molecules, our findings highlight a new, inducible mechanism of persistence, based on increased protection of ribosomes during stress, rather than slowdown of the metabolism.

In vitro characterization of RaiA in *E. coli*, has previously shown association with ribosomes during cold shock and stationary phase, but not during growth at 37°C, suggesting that binding of RaiA is prompted by stress (Agafonov et al., 2001). Based on crystal structures, RaiA was suggested to arrest translation (Vila-Sanjurjo et al., 2004). However, our results in *V. cholerae* do not support such a role, since there is no impact of RaiA overexpression on growth, and rather point to a protective effect of RaiA under ribosomal stress caused by AG treatment.

Alternatively, since RaiA is able to stabilize the 70S ribosome monomers against dissociation *in vitro* (Agafonov et al., 1999; Di Pietro et al., 2013), it was proposed to constitute a pool of inactive 70S ribosomes preserved from degradation in bacteria (Giuliodori, 2016; Giuliodori et al., 2007; Gualerzi et al., 2011).

*In vivo* effects of RaiA on bacterial phenotypes are less well described. A protective effect of RaiA during stress, such as starvation, was previously observed in the Gram-positive species *Mycobacterium tuberculosis* (Li et al., 2018) and *Lactococcus lactis* (Puri et al., 2014). Yet, the mechanisms remained enigmatic. Our results show a role of RaiA on survival to antibiotic stress in Gram-negative pathogens. One known mechanism of protection of non-translating ribosomes is ribosome hibernation. The ribosome hibernation factors, Rmf (ribosome modulation factor) and Hpf (hibernation promoting factor) (Maki et al., 2000), dimerize 70S ribosomes (monosomes) into 100S hibernating ribosome dimers. The importance of ribosome hibernation in stress survival is well established in various bacteria (McKay and Portnoy, 2015; Tkachenko et al., 2017), as 100S dimers are less susceptible to degradation by RNases (Prossliner et al., 2018; Wada et al., 2000; Yamagishi et al., 1993). Ribosome hibernation factors were even proposed as potential new targets for antibiotics (Matzov et al., 2019). However, in some cases, 70S particles appear to be more robust during heating than 100S dimers which dissociate into 30S and 50S subunits more rapidly (Niven, 2004).

RaiA was previously identified as bound to the ribosome together with Rmf and Hpf, but its role in relation with ribosome hibernation is unclear. In *V. cholerae*, RaiA shows a synergistic effect with Hpf for survival to starvation (Sabharwal et al., 2015), and we observe a synthetic lethal phenotype for the deletion of *raiA* and *hpf*. Despite such apparent synergy with hibernation factors, RaiA was shown to inactivate 70S ribosomes without forming 100S dimers (Polikanov et al., 2012; Ueta et al., 2005). RaiA thus appears to act in a process different than ribosome hibernation, maybe by blocking breaking down of ribosomes into 30S and 50S subunits by ribosome recycling factors (Rrf/EF-G) (Agafonov et al., 1999; Janosi et al., 1996).

In *E. coli*, RaiA can even prevent 100S dimer formation by Hpf and Rmf (Maki et al., 2000; Ueta et al., 2005). According to cryo-EM data, RaiA can compete with Rmf for ribosome binding, hence shifting the ribosome content from a Rmf-mediated dimeric inactive form to a RaiA-bound monomeric inactive form (Franken et al., 2017).

While RaiA, Hpf and Rmf are rapidly released from ribosomes when normal growth conditions are restored (Agafonov et al., 2001; Maki et al., 2000), ribosome reactivation necessitates dissociation of hibernating 100S ribosome dimers into monomers by HflX and other factors (Basu and Yap, 2017), whereas no dissociation factor is needed for the reactivation of the RaiA inactivated 70S ribosome. There may thus be an interplay and equilibrium between hibernating 100S-Hpf/Rmf and “sleeping” 70S-RaiA forms.

Such synergy or antagonism between inactive 70S and 100S ribosome pools can however depend on the nature of the stress and the bacterial species. We show here that persistence to aminoglycosides is better achieved in the presence of RaiA (70S), rather than hibernation factors (100S) in *V. cholerae*. Based on this, we propose that increased production of RaiA (e.g. upon stress) leads to preservation of ribosomes in a pool of inactive and intact 70S “sleeping” ribosomes. In that scenario, bacteria can keep growing in the absence of the antibiotic and avoid death upon antibiotic treatment, not by entering a dormant state, but by inducing protection of ribosomes. We propose that, in contrast to hibernating ribosomes, such sleeping ribosomes can be rapidly reactivated upon stress relief through spontaneous dissociation of RaiA, conferring an advantage for stress survival.

In line with this, a recent study showed that the greater the ribosome content of the cell, the faster persister cells resuscitate (Kim et al., 2018). In parallel, protein synthesis was shown to be necessary for persister cell formation in *V. cholerae* (Paranjape and Shashidhar, 2019), consistent with the fact that we see no effect of RaiA overexpression on growth rates. Our results do not exclude however another role for RaiA on translating ribosomes.

Increased RaiA levels thus allow higher persistence, but what is controlling RaiA expression? RaiA is expressed in stationary phase and during cold shock (Agafonov et al., 1999; Agafonov et al., 2001; Maki et al., 2000; Sabharwal et al., 2015). Other known factors are the heat shock in *V. cholerae* (Slamti et al., 2007), stringent response (Prossliner et al., 2018), envelope stress and the carbon catabolite response (Manneh-Roussel et al., 2018; Shimada et al., 2013). We additionally show that *raiA* expression is linked to iron levels and responds to Fur regulation. Interestingly, iron associates with ribosomes (Bray et al., 2018) and a link between iron and modulation of ribosome function during stress has been described (Zinskie et al., 2018). Iron control of RaiA by may thus allow protection of ribosomes from iron related damage. Altogether, RaiA appears to be part of the bacterial response to environmental stress.

Results of the present study constitute a link between bacterial signaling, ribosome protection and persistence. Moreover, since RaiA action appears to be conserved in various pathogens (Gram-negatives and positives, such as *S. aureus* and *M. tuberculosis*), it may be one factor involved in the failure of treatment in persistent infections. It would be interesting to ask now whether RaiA can be used as an early indicator of persistence, which would allow isolation of persisters within a heterogeneous population and further studies using single cell approaches.

## Acknowledgements

We are thankful to Micheline Fromont-Racine for her valuable help with the experiments for ribosome content analysis by sucrose density gradients. We thank Ivan Matic, Wei-Lin Su and Sébastien Fleurier for helpful discussions and Ivan Matic for critical reading of the manuscript. We also thank Sebastian Aguilar Pierlé for advice for Neo-Cy5 uptake experiments. This work was supported by the Institut Pasteur, the Centre National de la Recherche Scientifique (CNRS-UMR 3525), the Fondation pour la Recherche Médicale (FRM Grant No. DBF20160635736), ANR Unibac (ANR-17-CE13-0010-01) and Institut Pasteur grant PTR 245-19.

## Methods

**Mutant strains and plasmids** are described in table S2.

### Persistence tests

Persistence tests were performed on early exponential phase cultures. In order to clear the culture from previously non-growing cells that could potentially be present from the stationary phase inoculum, we performed a two-step dilution protocol, before antibiotic treatment. Overnight *V. cholerae* cultures were first diluted 1000x in 4 ml fresh Mueller-Hinton (MH) medium, without indole (with the antibiotic allowing to maintain the plasmid, when the strain carried a plasmid) and incubated at 37°C with shaking. When the OD620 nm reached 0,2, cultures were diluted 1000x a second time, in order to clear them from non-growing cells, in Erlenmeyers containing 25 ml fresh MH medium, without or with indole at 350 µM (with the antibiotic allowing to maintain the plasmid, when the strain carried a plasmid), and were allowed to grow at 37°C. When cultures reached an OD_620nm_ between 0,25 and 0,3 (early exponential phase), appropriate dilutions were plated on MH plates to determine the total number of CFUs in time zero untreated cultures. Note that for *V. cholerae*, it was important to treat cultures at the precise OD 0,25-0,3, as persistence levels seem to be sensitive to growth phase in *V. cholerae*, where they decline in stationary phase, and because we wanted to avoid any stationary phase protein expression such as *raiA* or *rpoS* at later growth. 5 ml of cultures were collected into 50 ml Falcon tubes and treated with lethal doses of desired antibiotics (5-10 times the MIC: tobramycin 10 µg/ml, gentamycin 5 µg/ml, carbenicillin 100 µg/ml) for 20 hours at 37°C with shaking in order to guarantee oxygenation. Appropriate dilutions were then plated on MH agar without antibiotics and proportion of growing CFUs were calculated by doing a ration with total CFUs at time zero. Experiments were performed 3 to 6 times.

**Quantification of fluorescent neomycin uptake** was performed as described (Pierlé et al, manuscript in revision). Briefly, overnight cultures were diluted 100-fold in M9 minimal medium supplemented with 0.4% glucose. When the bacterial strains reached an OD_620nm_ of 0.25, they were incubated with 0.4 µM of Cy5 labeled Neomycin for 15 minutes at 37°C. 10 µl of the incubated culture were then used for flow cytometry, diluting them in 250 μl of PBS before reading fluorescence. WT *V. cholerae*, was incubated simultaneously without Neo-Cy5 as a negative control. Flow cytometry experiments were performed as described (Baharoglu et al., 2010) and repeated at least 3 times. For each experiment, 100000 events were counted on the Miltenyi MACSquant device.

#### MIC determination using *etests*

Stationary phase cultures were diluted 20 times in PBS, and 300 µL were plated on MH plates and dried for 10 minutes. *etests* (Biomérieux) were placed on the plates and incubated overnight at 37°C.

#### RNA-seq

Overnight cultures of the O1 biovar El Tor N16961 *hapR+ V. cholerae* strain were diluted 100x and grown in triplicate in MH medium until an OD_620nm_ 0.4 with or without 350 µM indole. Sample collection, total RNA was extraction, library preparation, sequencing and analysis were performed as previously described (Krin et al., 2018).

#### *raiA* qRT-PCR

Total RNA was extracted and purified after 6 hours cultures in MH in presence or absence of indole, as previously described (Krin et al., 2018). Reverse transcription (RT) was performed on 100 ng total RNA using SuperScript^®^ III First-Strand Synthesis System for RT-PCR (Invitrogen). Quantitative PCR was performed on 2 µl RT sample diluted 10-fold using SYBR Green PCR Master Mix (APPLIED) and QuantStudio 6. Quantification was performed using standard range.

#### Quantification of *raiA* expression by fluorescent flow cytometry using a *gfp* fusion

*gfp* was amplified by PCR using primers carrying the *raiA* promoter region and cloned into pTOPO-TA cloning vector. The PraiA-gfp fragment was then extracted using EcoRI and cloned into the low copy plasmid pSC101 (1 to 5 copies per cell). The plasmid was introduced into desired strains, and fluorescence was measured by on specified conditions, by counting 100000 cells on the Miltenyi MACSquant device.

#### Growth curves

Overnight cultures were diluted 100x in fresh medium, on 96 well plates. Each well contained 200 µl. Plates were incubated with shaking on TECAN device at 37°C, OD 620 nm was measured every 15 minutes.

### Preparation of cell lysate for the analysis of ribosome content

The protocol was adapted from (Qin and Fredrick, 2013). Since we used stationary phase cultures instead of exponential phase, presence of polysomes is not expected. 10 ml of 20-hour cultures were centrifuged in ice cold 50ml Falcon tubes for 15min at 5000 rpm at 4°C. Pellets were resuspended in 500µl lysis buffer (10mM Tris-HCl, pH 8, 10mM MgCl2, Lysozyme 1mg/ml, protease inhibitor), transferred in ice cold 1,5ml tubes and incubated with 12 µl Ribolock RNase inhibitor (Thermo scientific) and DNaseI (5U/ml) at 4°C for 15min. Cell lysis was performed through 3 cycles of flash-freezing in dry ice and thawing in a water bath at 4°C. 15µl of 10% sodium deoxycholate were added and cell lysate was obtained after centrifugation at 10 000 rpm for 10 minutes at 4°C. The pellet containing cell debris was discarded. Lysate was kept at −80°C until sucrose gradient ultracentifugation.

### Sucrose gradient

10-50% sucrose gradient tubes (Beckman ULTRA CLEAR) were prepared. 2U of OD260 of each cell extracts were deposited on sucrose gradient tubes. Ultracentrifugation was performed at 39 000 rpm at 4°C for 2h45. Fractions were collected using a pump coupled to a spectrometer at OD 260nm, and plotted as a function of time (half seconds).

## Supplemental information

**Table S1. Differentially expressed genes upon indole treatment**.

**Table S2. Strains and plasmids**.

**Supplementary text, methods and figure legends**.

**Quantification of extracellular indole concentrations (Figure S1)**. In order to address whether indole secretion is increased in the presence of sub-MIC TOB in *V. cholerae*, we measured extracellular indole concentrations of bacterial cultures grown overnight in rich media with and without antibiotics using the Kovacs reagent (Saint-Ruf et al., 2014). We used two different media: the Mueller-Hinton (MH) that we classically use for experiments involving sub-MIC antibiotics, and an additional defined rich medium (Teknova EZ rich defined medium), hereafter called rich MOPS. The MIC of TOB was measured for *V. cholerae* using *etests* and is 1-1.2 µg/ml in MH, and 0.75 µg/ml in rich MOPS. Unexpectedly, we found no indole production in MH medium in any condition (not shown). Indole is produced in rich medium (DOI: 10.1006/jmbi.1999.3462) and MH might not be rich enough to allow its production from tryptophan. In rich MOPS (as well as in LB, not shown), we found that sub-MIC TOB (from 20% of the MIC) significantly increases indole secretion in WT *V. cholerae*. The following experiments were thus conducted in MH, as it allows to study the impact of defined indole concentrations added to the media.

**Figure S1. Indole is produced during growth in sub-MIC tobramycin and improves growth of *V. cholerae* in presence of antibiotics. A**. Measure of extracellular indole concentrations of bacterial cultures grown overnight in rich medium MOPS (Teknova EZ rich defined medium) with and without antibiotics using the Kovacs reagent (Saint-Ruf et al., 2014) *ΔtnaA* strain was used as negative control. **BCD**. Growth is measured on a TECAN plate reader by measuring OD at 620nm (y axis), for the time indicated by the x axis. IND: indole 350µM. TOB: tobramycin sub-MIC (0,6 µg/ml). CM: chloramphenicol sub-MIC (0,4µg/ml). Indole has no effect on growth in the absence of antibiotics and improves growth in the presence of sub-MIC TOB and CM, thus independently of the family of antibiotics.

**Figure S2. Persistence and the effect of indole and RaiA. A**. Kinetics of survival of WT *V. cholerae* to tobramycin 10µg/ml in MH. Time zero corresponds to the total number of CFUs before addition of antibiotics, to an early exponential phase culture (OD_620nm_ 0,25-0,3), as described in the methods section. The proportion of surviving CFUs is calculated after plating and counting growing colonies, and is represented for each time point. Each curve represents one replicate **B**. Persistence of *V. cholerae* WT and Δ*tnaA* mutants at exponential phase (in LB to allow indole production) after 20 hours treatment with specified antibiotics. **C**. Persistence of *E. coli* WT and *raiA* mutant, at late exponential phase (OD_620nm_ 0,5), after 20 hours treatment with 10µg/ml tobramycin (TOB) in MH. **D**. Persistence of *V*. cholerae WT and mutants at exponential phase in MH. The same persistence protocol as described above for A was used and serial dilutions are spotted on plates for estimation of survival.

**Figure S3. RaiA levels influence stationary phase 70S ribosome content relative to 50S+30S subunits in *V. cholerae* in presence of tobramycin**. Cellular extracts of 24 hours cultures in tobramycin (sub-MIC, 0,6µg/ml) were separated on 10-50% sucrose density gradient. Ribosomal RNA content was measured at OD 260nm using a spectrometer coupled to a pump and time on X axis represents samples from less dense (upper fragments, smaller complexes) to denser (bottom of the tube, heavier complexes). Cell debris eluting before 1100 half seconds are not shown. Graphs are normalized to total OD260nm=1 for each sample.

