## Supplemental figures and tables for "Sleeping ribosomes: bacterial signaling triggers RaiA mediated persistence to aminoglycosides"

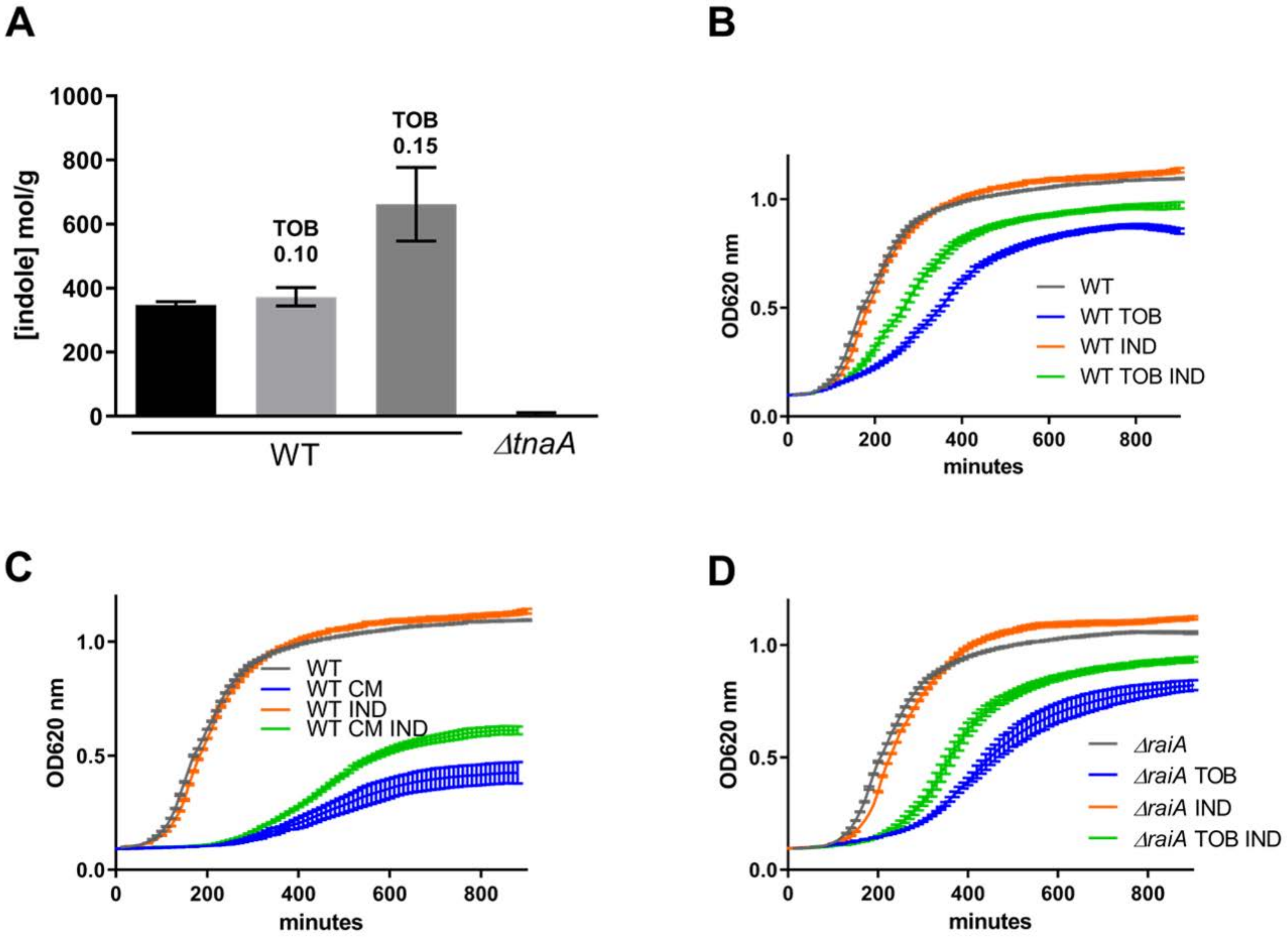
A
1000
800
600
400
200
0
[indole] mol/g
TOB
0.10
TOB
0.15
WT
ΔtnaA
B
OD620 nm
1.0
0.5
0.0
0
200
400
600
800
minutes
WT
WT TOB
WT IND
WT TOB IND
C
OD620 nm
1.0
0.5
0.0
0
200
400
600
800
minutes
WT
WT CM
WT IND
WT CM IND
D
OD620 nm
1.0
0.5
0.0
0
200
400
600
800
minutes
ΔraiA
ΔraiA TOB
ΔraiA IND
ΔraiA TOB IND

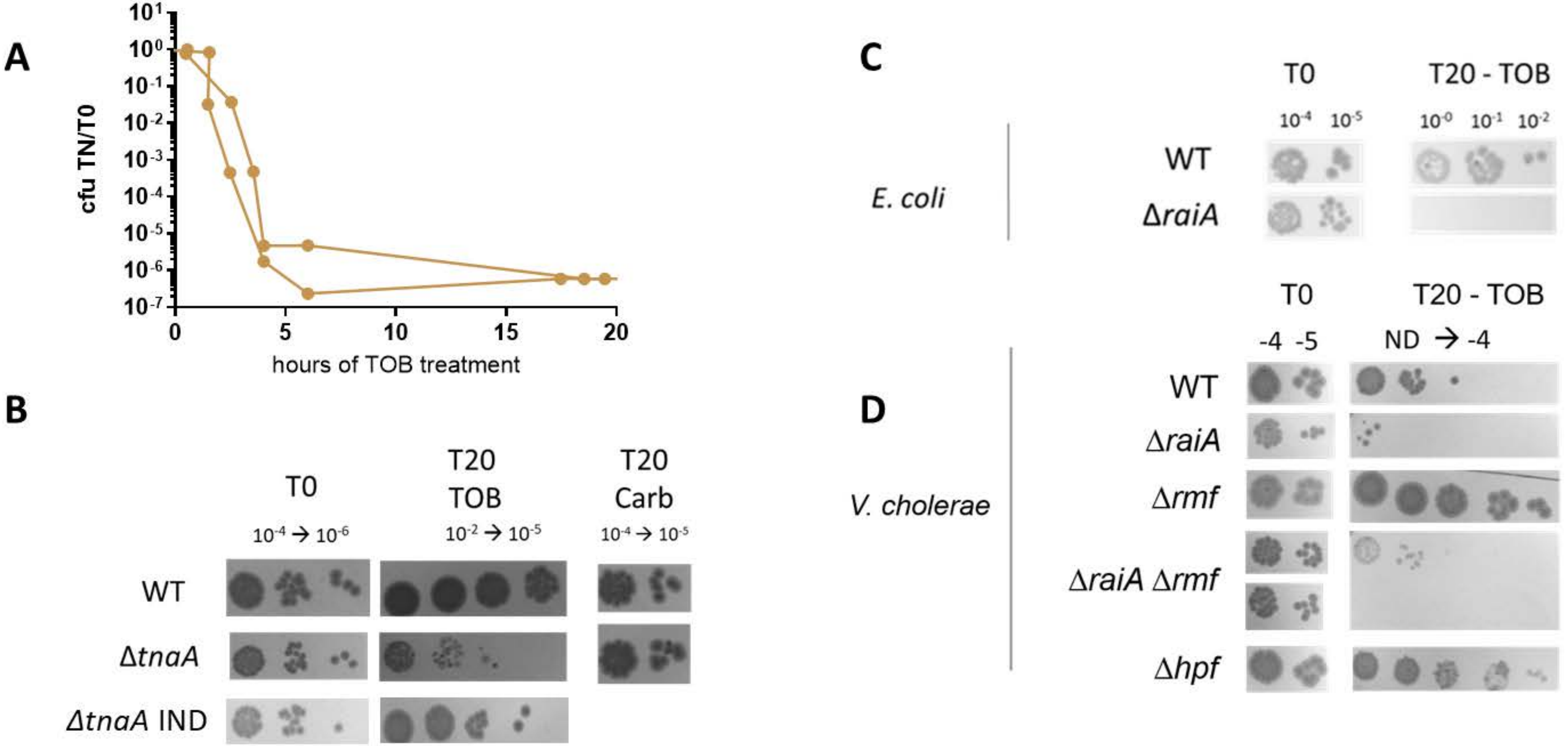

A
cfu TN/T0
hours of TOB treatment
B
T0
T20 TOB
T20 Carb
WT
ΔtnaA
ΔtnaA IND
C
T0
T20 - TOB
E. coli
WT
ΔraiA
D
T0
T20 - TOB
-4 -5
ND → -4
V. cholerae
WT
ΔraiA
Δrmf
ΔraiA Δrmf
Δhpf

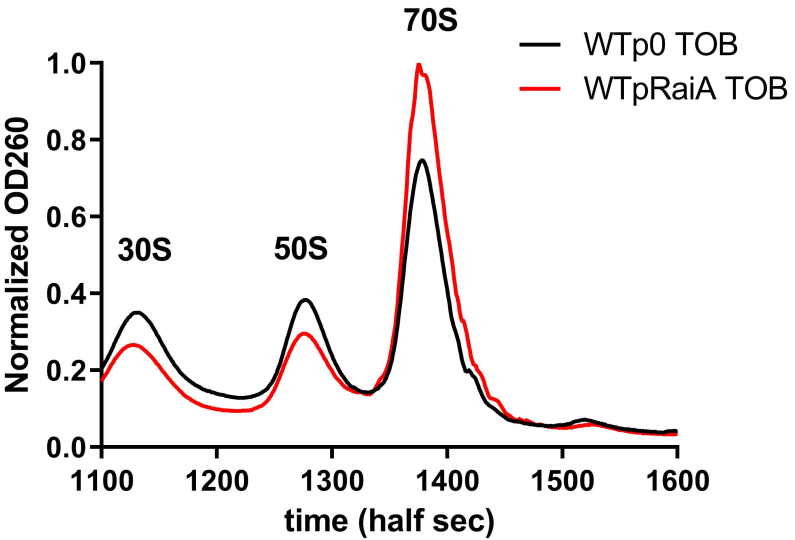
70S
WTp0 TOB
WTpRaiA TOB
1.0
0.8
0.6
0.4
0.2
0.0
Normalized OD260
30S
50S
1100
1200
1300
1400
1500
1600
time (half sec)

**Table S1. Genes differentially regulated by indole.**

| Id | norm MH-A | norm MH-B | norm MH-C | norm IND-A | norm IND-B | norm IND-C | **Fold change** | p value | adj p value |
|---|---|---|---|---|---|---|---|---|---|
| UP | | | | | | | | | |
| VC0706 | 3903 | 4103 | 3710 | 80171 | 68404 | 145086 | 20,1 | 8,0E-03 | 8,9E-03 |
| VC1484 | 120 | 111 | 142 | 570 | 232 | 501 | 3,1 | 2,6E-05 | 1,3E-01 |
| Motility- Chemotaxis | | | | | | | | | |
| VC0410 | 1779 | 1337 | 1477 | 3761 | 3910 | 3890 | **2,5** | 1,7E-14 | 8,0E-03 |
| VC1049 | 413 | 473 | 450 | 2398 | 2145 | 2322 | **4,9** | 1,1E-38 | 9,0E-03 |
| VC2370 | 316 | 270 | 299 | 1469 | 1694 | 1286 | **4,8** | 2,6E-35 | 1,0E-02 |
| VC0414 | 3748 | 3638 | 2441 | 7061 | 7532 | 6276 | **2,1** | 1,4E-06 | 1,9E-02 |
| VC0413 | 658 | 597 | 376 | 1239 | 1302 | 1053 | **2,1** | 1,2E-05 | 3,0E-02 |
| VC0411 | 1939 | 1418 | 1067 | 3411 | 3912 | 2746 | **2,2** | 1,6E-05 | 3,9E-02 |
| VCA0536 | 508 | 934 | 598 | 1748 | 2438 | 2062 | **2,9** | 6,5E-09 | 4,6E-02 |
| VCA0954 | 1750 | 2718 | 1721 | 3886 | 5756 | 4105 | **2,1** | 2,7E-04 | 5,9E-02 |
| VC0409 | 9371 | 6775 | 5433 | 26356 | 19176 | 30086 | **3,3** | 3,7E-09 | 6,3E-02 |
| VC0827 | 1152 | 1207 | 1001 | 3000 | 7845 | 2896 | **3,6** | 4,4E-07 | 1,4E-01 |
| VC0826 | 1967 | 1966 | 1336 | 5009 | 13957 | 5414 | **4,0** | 4,7E-08 | 1,5E-01 |
| VC1763 | 428 | 1057 | 267 | 1554 | 1847 | 1495 | **2,5** | 3,3E-03 | 2,0E-01 |
| VC0820 | 369 | 623 | 278 | 1133 | 751 | 2000 | **2,6** | 1,5E-03 | 2,2E-01 |
| Respiration - energy | | | | | | | | | |
| VC1115 | 663 | 546 | 508 | 6006 | 6126 | 5482 | **9,8** | 6,1E-84 | 7,9E-03 |
| VC0966 | 7486 | 7229 | 6500 | 15713 | 18048 | 21828 | **2,6** | 2,3E-14 | 1,1E-02 |
| VC2371 | 163 | 157 | 107 | 1820 | 1907 | 1910 | **12,1** | 6,9E-67 | 1,3E-02 |
| VC0477 | 4301 | 4841 | 4095 | 27291 | 30017 | 20290 | **5,6** | 4,2E-39 | 1,5E-02 |
| VCA0511 | 174 | 249 | 197 | 1325 | 1518 | 1124 | **6,0** | 4,6E-35 | 1,8E-02 |
| VCA0665 | 329 | 253 | 320 | 9542 | 8962 | 8507 | **27,0** | 5,2E-122 | 1,8E-02 |
| VCA0665 | 329 | 253 | 320 | 9542 | 8962 | 8507 | **27,0** | 5,2E-122 | 1,8E-02 |
| VC1951 | 347 | 334 | 242 | 5920 | 6998 | 5337 | **17,9** | 1,2E-87 | 2,0E-02 |
| VC1114 | 430 | 337 | 365 | 1584 | 1201 | 1615 | **3,7** | 1,7E-17 | 2,3E-02 |
| VC0356 | 583 | 766 | 592 | 1725 | 2375 | 1675 | **2,9** | 7,9E-12 | 2,4E-02 |
| VC2145 | 931 | 952 | 989 | 7482 | 9974 | 5936 | **7,6** | 1,1E-42 | 2,4E-02 |
| VCA0563 | 3532 | 3857 | 3297 | 10041 | 12683 | 8503 | **2,8** | 3,1E-11 | 2,7E-02 |
| VC2368 | 8558 | 11124 | 8314 | 25789 | 18044 | 26995 | **2,4** | 4,1E-08 | 2,9E-02 |
| VC0278 | 512 | 493 | 641 | 1311 | 1930 | 1207 | **2,6** | 1,2E-08 | 3,1E-02 |
| VC0485 | 8659 | 11085 | 11206 | 35439 | 45566 | 26147 | **3,3** | 4,0E-13 | 3,2E-02 |
| VCA0564 | 4137 | 4464 | 3746 | 10730 | 12961 | 8158 | **2,5** | 8,6E-08 | 3,4E-02 |
| VC2656 | 941 | 1355 | 964 | 33710 | 22990 | 28100 | **22,5** | 2,0E-76 | 4,1E-02 |
| VC2646 | 2110 | 1858 | 1643 | 4701 | 6486 | 4176 | **2,6** | 1,7E-07 | 4,6E-02 |
| VC1687 | 61 | 69 | 112 | 1751 | 1301 | 1331 | **15,5** | 6,7E-48 | 5,0E-02 |
| VC1866 | 7913 | 12676 | 7185 | 81168 | 101349 | 63711 | **7,9** | 6,8E-31 | 5,3E-02 |
| VC2657 | 195 | 285 | 147 | 10748 | 7717 | 8506 | **33,9** | 1,1E-70 | 7,0E-02 |

**Table S1. Genes differentially regulated by indole.**

| | | | | | | | | | |
|---|---|---|---|---|---|---|---|---|---|
| VC1516 | 769 | 1005 | 666 | 2063 | 981 | 2137 | **2,0** | 2,6E-03 | 7,9E-02 |
| VC2699 | 895 | 1117 | 1191 | 5749 | 3169 | 5283 | **4,0** | 1,1E-09 | 9,3E-02 |
| VCA0675 | 293 | 225 | 186 | 1644 | 816 | 2194 | **5,7** | 1,1E-14 | 9,5E-02 |
| VC0965 | 7652 | 17113 | 10909 | 25906 | 35487 | 21426 | **2,2** | 1,2E-03 | 9,7E-02 |
| VC1513 | 979 | 1330 | 598 | 4220 | 2083 | 4053 | **3,2** | 9,9E-07 | 1,0E-01 |
| VC2447 | 25911 | 36038 | 27345 | 51499 | 105140 | 37661 | **2,0** | 7,0E-03 | 1,3E-01 |
| VCA0680 | 583 | 386 | 577 | 2435 | 870 | 2782 | **3,5** | 1,5E-06 | 1,3E-01 |
| VCA0679 | 626 | 383 | 350 | 2412 | 805 | 2866 | **3,8** | 9,4E-07 | 1,6E-01 |
| VCA0678 | 1612 | 1152 | 1434 | 5851 | 1605 | 7354 | **3,0** | 1,3E-04 | 1,9E-01 |
| Sugar | | | | | | | | | |
| VC2000 | 33134 | 38523 | 32617 | 94393 | 96880 | 78059 | **2,5** | 2,7E-15 | 9,0E-03 |
| VC0374 | 3016 | 2691 | 2733 | 9064 | 9875 | 7214 | **3,0** | 2,5E-18 | 1,2E-02 |
| VC1973 | 609 | 640 | 690 | 6655 | 6173 | 5216 | **8,8** | 4,0E-64 | 1,3E-02 |
| VC1972 | 229 | 204 | 255 | 2990 | 2618 | 2426 | **10,7** | 3,5E-58 | 2,0E-02 |
| VC1971 | 280 | 208 | 250 | 3269 | 2917 | 2594 | **11,0** | 1,4E-58 | 2,2E-02 |
| VC0487 | 2056 | 1941 | 1773 | 12473 | 9768 | 9035 | **5,1** | 6,3E-27 | 2,8E-02 |
| VCA0011 | 487 | 642 | 554 | 2564 | 2618 | 2403 | **4,3** | 9,8E-20 | 2,9E-02 |
| VC2689 | 3011 | 2941 | 4636 | 23493 | 19880 | 18705 | **5,4** | 1,3E-24 | 3,9E-02 |
| VC2001 | 1039 | 618 | 617 | 2290 | 3235 | 1950 | **3,1** | 8,4E-09 | 5,9E-02 |
| VCA0516 | 258 | 256 | 239 | 1777 | 2151 | 1542 | **6,5** | 7,7E-22 | 6,0E-02 |
| VCA0013 | 493 | 1132 | 476 | 2028 | 1287 | 1755 | **2,2** | 7,3E-03 | 1,7E-01 |
| RRR | | | | | | | | | |
| VCA0521 | 1439 | 1451 | 801 | 3686 | 3125 | 3167 | **2,6** | 3,1E-07 | 4,4E-02 |
| VCA0520 | 694 | 723 | 482 | 1449 | 1115 | 1411 | **2,0** | 3,1E-05 | 2,9E-02 |
| VC2386 | 4774 | 8304 | 2645 | 8736 | 6458 | 24643 | **2,2** | 3,3E-02 | 3,6E-01 |
| Metabol | | | | | | | | | |
| VC0336 | 7124 | 8934 | 9210 | 19907 | 24075 | 16926 | **2,4** | 2,8E-09 | 1,8E-02 |
| VC0026 | 268 | 258 | 311 | 1582 | 1606 | 1262 | **5,0** | 1,6E-27 | 2,1E-02 |
| VCA0235 | 3581 | 3566 | 3392 | 8268 | 5703 | 8159 | **2,0** | 5,8E-05 | 3,3E-02 |
| VCA0875 | 2063 | 2616 | 1307 | 5759 | 4746 | 4221 | **2,4** | 7,0E-06 | 4,4E-02 |
| VC2698 | 2245 | 3759 | 3380 | 7765 | 5269 | 7196 | **2,1** | 6,6E-04 | 5,7E-02 |
| VC2373 | 1415 | 690 | 1058 | 2458 | 3824 | 2252 | **2,5** | 6,8E-05 | 8,7E-02 |
| VC2361 | 4108 | 2170 | 3027 | 19721 | 15393 | 32664 | **6,1** | 1,1E-13 | 1,3E-01 |
| Nucleotide | | | | | | | | | |
| VCA0564 | 4137 | 4464 | 3746 | 10730 | 12961 | 8158 | **2,5** | 8,6E-08 | 3,4E-02 |
| VC1034 | 470 | 448 | 562 | 1528 | 1273 | 1225 | **2,7** | 5,9E-13 | 1,2E-02 |
| regulators | | | | | | | | | |
| VC0486 | 1203 | 1242 | 753 | 11275 | 8624 | 10176 | **8,5** | 1,0E-37 | 3,6E-02 |
| VC1719 | 1362 | 981 | 674 | 5308 | 5146 | 6057 | **5,1** | 4,1E-20 | 4,3E-02 |
| VCA0562 | 824 | 717 | 348 | 1268 | 1261 | 1628 | **2,1** | 9,6E-04 | 7,1E-02 |
| VC1919 | 12704 | 12883 | 7383 | 28846 | 15905 | 39769 | **2,4** | 1,1E-03 | 1,2E-01 |
| VC2366 | 483 | 775 | 287 | 1133 | 1549 | 946 | **2,2** | 3,7E-03 | 1,2E-01 |

**Table S1. Genes differentially regulated by indole.**

| | | | | | | | | | |
|---|---|---|---|---|---|---|---|---|---|
| csrD | 58131 | 44933 | 26252 | 102802 | 52169 | 157915 | **2,2** | 1,0E-02 | 1,8E-01 |
| Membrane | | | | | | | | | |
| VC1315 | 54 | 49 | 49 | 1059 | 987 | 1071 | **18,6** | 6,1E-84 | 1,0E-08 |
| VC2484 | 2791 | 2555 | 2045 | 8396 | 7987 | 7622 | **3,2** | 4,9E-20 | 1,1E-02 |
| VC2485 | 2791 | 2555 | 2045 | 8396 | 7987 | 7622 | **3,2** | 4,9E-21 | 1,1E-02 |
| VCA0591 | 531 | 462 | 342 | 1184 | 1285 | 1183 | **2,7** | 9,1E-12 | 1,6E-02 |
| VC0976 | 1443 | 1713 | 1027 | 3220 | 2738 | 3167 | **2,1** | 1,4E-05 | 3,1E-02 |
| VC2149 | 145 | 110 | 144 | 2554 | 1821 | 2827 | **15,4** | 6,3E-47 | 5,7E-02 |
| VC0654 | 606 | 731 | 365 | 2927 | 5686 | 3163 | **5,9** | 8,2E-14 | 1,1E-01 |
| VCA0628 | 680 | 1842 | 822 | 2839 | 1462 | 4368 | **2,2** | 1,7E-02 | 2,9E-01 |
| Others | | | | | | | | | |
| VC0652 | 74 | 71 | 98 | 1343 | 1362 | 1262 | **15,1** | 3,0E-86 | 1,0E-08 |
| VC0428 | 523 | 576 | 695 | 4516 | 4519 | 4760 | **7,3** | 1,6E-49 | 1,6E-02 |
| VC1892 | 819 | 567 | 543 | 1182 | 2075 | 1046 | **2,1** | 5,6E-04 | 7,2E-02 |
| VC0037 | 801 | 555 | 621 | 1749 | 1301 | 2937 | **2,8** | 6,0E-06 | 8,9E-02 |
| VC1765 | 1538 | 4879 | 1100 | 7066 | 10157 | 6196 | **2,6** | 3,1E-03 | 2,9E-01 |
| Hyp | | | | | | | | | |
| VCA0718 | 329 | 301 | 343 | 1989 | 1854 | 1881 | **5,7** | 6,9E-63 | 1,0E-08 |
| VC1865 | 741 | 620 | 752 | 15085 | 13445 | 13011 | **18,5** | 7,9E-143 | 7,1E-03 |
| VC1871 | 630 | 784 | 729 | 5093 | 6873 | 5480 | **7,7** | 7,7E-58 | 1,3E-02 |
| VC1077 | 1496 | 1106 | 1197 | 2628 | 2923 | 2918 | **2,2** | 1,1E-08 | 1,4E-02 |
| VCA0236 | 299 | 343 | 418 | 1039 | 1012 | 978 | **2,8** | 2,1E-12 | 1,5E-02 |
| VCA1013 | 41 | 54 | 72 | 1544 | 916 | 1414 | **19,5** | 9,5E-54 | 5,0E-02 |
| VCA0741 | 8417 | 7150 | 6445 | 35536 | 27576 | 52687 | **4,8** | 2,1E-17 | 5,5E-02 |
| VCA0919 | 1107 | 806 | 959 | 3604 | 2763 | 4638 | **3,6** | 4,7E-11 | 5,5E-02 |
| VC0871 | 209 | 193 | 259 | 1657 | 1019 | 2475 | **6,8** | 3,3E-19 | 8,0E-02 |
| VC1853 | 2696 | 2439 | 1227 | 5889 | 3740 | 7203 | **2,4** | 3,1E-04 | 9,6E-02 |
| VC2035 | 272 | 247 | 652 | 883 | 1154 | 1004 | **2,4** | 4,9E-04 | 1,1E-01 |
| VCA0367 | 870 | 2025 | 677 | 2495 | 2608 | 3104 | **2,1** | 6,6E-03 | 1,5E-01 |
| VCA0547 | 1134 | 821 | 2003 | 3510 | 1518 | 3735 | **2,0** | 2,3E-02 | 1,9E-01 |
| VC1770 | 338 | 770 | 245 | 1049 | 1704 | 933 | **2,4** | 5,3E-03 | 2,1E-01 |
| VC2221 | 97 | 191 | 99 | 3971 | 1113 | 3984 | **15,1** | 9,8E-21 | 2,4E-01 |
| VCA0283 | 721 | 1944 | 390 | 2479 | 2047 | 3333 | **2,2** | 1,7E-02 | 2,6E-01 |
| VC1764 | 769 | 2351 | 507 | 4476 | 4878 | 3997 | **3,0** | 5,2E-04 | 2,8E-01 |
| DOWN | | | | | | | | | |
| Electron transfer - Ion | | | | | | | | | |
| VC2415 | 6690 | 8047 | 8130 | 1766 | 1466 | 1837 | 0,2 | 3,9E-40 | 6,4E-38 |
| VCA0228 | 703 | 914 | 818 | 60 | 32 | 108 | 0,1 | 8,9E-25 | 7,7E-23 |
| VC1265 | 1746 | 1306 | 1376 | 237 | 112 | 243 | 0,2 | 8,6E-24 | 6,8E-22 |
| VCA0227 | 2696 | 3888 | 2205 | 333 | 155 | 375 | 0,1 | 4,4E-22 | 3,3E-20 |
| VCA0230 | 883 | 1120 | 957 | 128 | 53 | 173 | 0,1 | 5,5E-19 | 3,1E-17 |
| VCA0229 | 545 | 636 | 685 | 68 | 15 | 72 | 0,1 | 6,8E-19 | 3,7E-17 |
| VC2559 | 1531 | 1298 | 1238 | 326 | 158 | 308 | 0,2 | 1,3E-16 | 6,1E-15 |

**Table S1. Genes differentially regulated by indole.**

| | | | | | | | | | |
|---|---|---|---|---|---|---|---|---|---|
| VC2414 | 20657 | 25736 | 21516 | 6744 | 3908 | 6762 | 0,3 | 2,5E-16 | 1,2E-14 |
| VC1434 | 5799 | 4818 | 6784 | 1544 | 2021 | 1387 | 0,3 | 1,6E-15 | 6,9E-14 |
| VC1544 | 860 | 841 | 702 | 211 | 92 | 200 | 0,2 | 5,8E-15 | 2,3E-13 |
| VC1266 | 992 | 730 | 899 | 180 | 81 | 218 | 0,2 | 2,8E-14 | 1,1E-12 |
| VC1548 | 828 | 977 | 762 | 200 | 71 | 185 | 0,2 | 3,5E-14 | 1,3E-12 |
| VC1168 | 2427 | 1657 | 3276 | 600 | 422 | 657 | 0,2 | 4,4E-14 | 1,6E-12 |
| VC2560 | 938 | 837 | 656 | 178 | 57 | 151 | 0,2 | 1,6E-13 | 5,6E-12 |
| VC0575 | 3481 | 4499 | 4787 | 1741 | 1310 | 1531 | 0,4 | 1,7E-13 | 5,9E-12 |
| VC0540 | 1091 | 965 | 938 | 257 | 204 | 349 | 0,3 | 1,9E-13 | 6,4E-12 |
| VC1890 | 4135 | 4112 | 4697 | 2193 | 1882 | 2160 | 0,5 | 2,8E-13 | 9,2E-12 |
| VC1267 | 1006 | 697 | 951 | 208 | 95 | 240 | 0,2 | 1,2E-12 | 3,6E-11 |
| VC1625 | 6073 | 5424 | 5715 | 1808 | 1036 | 2011 | 0,3 | 1,3E-12 | 3,8E-11 |
| VC1547 | 1186 | 1193 | 907 | 270 | 101 | 276 | 0,2 | 2,0E-12 | 5,9E-11 |
| VC1623 | 2356 | 1972 | 1754 | 913 | 895 | 853 | 0,4 | 2,2E-12 | 6,6E-11 |
| VC0608 | 1724 | 3895 | 2700 | 607 | 375 | 613 | 0,2 | 2,0E-11 | 5,5E-10 |
| VC1543 | 1647 | 1544 | 1204 | 468 | 228 | 484 | 0,3 | 2,7E-11 | 7,1E-10 |
| VC1264 | 3449 | 2931 | 2531 | 663 | 257 | 830 | 0,2 | 3,0E-11 | 7,8E-10 |
| VC0541 | 2020 | 1993 | 2192 | 759 | 645 | 969 | 0,4 | 6,9E-11 | 1,7E-09 |
| VC0627 | 1608 | 1342 | 1589 | 479 | 458 | 718 | 0,4 | 3,9E-10 | 8,7E-09 |
| VC2413 | 11184 | 12876 | 15480 | 4976 | 2760 | 5137 | 0,3 | 4,8E-10 | 1,0E-08 |
| VC0574 | 4963 | 6302 | 6116 | 2629 | 1626 | 2374 | 0,4 | 5,1E-10 | 1,1E-08 |
| VCA0779 | 1014 | 1043 | 1299 | 573 | 473 | 544 | 0,5 | 6,7E-10 | 1,4E-08 |
| VC1425 | 18728 | 18568 | 11818 | 6031 | 6511 | 6308 | 0,4 | 2,6E-09 | 4,9E-08 |
| VC1190 | 3535 | 3491 | 3519 | 1105 | 534 | 1290 | 0,3 | 4,7E-09 | 8,7E-08 |
| VC0539 | 816 | 759 | 978 | 299 | 125 | 291 | 0,3 | 5,5E-09 | 1,0E-07 |
| VC2031 | 1195 | 956 | 1683 | 516 | 398 | 518 | 0,4 | 6,3E-09 | 1,1E-07 |
| VC0538 | 2463 | 3016 | 3026 | 969 | 418 | 946 | 0,3 | 6,6E-09 | 1,2E-07 |
| VC2044 | 4468 | 3773 | 4970 | 2081 | 1443 | 2069 | 0,4 | 6,9E-09 | 1,2E-07 |
| VC1624 | 2450 | 2405 | 2567 | 1013 | 552 | 1133 | 0,4 | 3,1E-08 | 5,0E-07 |
| VC0384 | 2708 | 2991 | 1808 | 989 | 444 | 818 | 0,3 | 5,1E-08 | 8,1E-07 |
| VC1350 | 1809 | 1972 | 2095 | 700 | 899 | 926 | 0,4 | 6,0E-08 | 9,2E-07 |
| VC2099 | 4134 | 4299 | 6134 | 2367 | 2308 | 2392 | 0,5 | 1,8E-07 | 2,5E-06 |
| VC1299 | 2194 | 1529 | 1452 | 766 | 807 | 883 | 0,5 | 3,3E-06 | 3,5E-05 |
| VC2389 | 12335 | 10446 | 14700 | 5767 | 2538 | 5581 | 0,4 | 3,7E-06 | 3,8E-05 |
| VC0168 | 4710 | 4663 | 3918 | 2115 | 1538 | 2593 | 0,5 | 5,0E-06 | 5,0E-05 |
| VC1439 | 3658 | 4680 | 4161 | 2062 | 1186 | 2295 | 0,5 | 5,5E-06 | 5,5E-05 |
| VC0385 | 2632 | 2744 | 1443 | 910 | 465 | 974 | 0,4 | 6,3E-06 | 6,3E-05 |
| VC0386 | 1716 | 1717 | 944 | 651 | 435 | 748 | 0,4 | 2,9E-05 | 2,5E-04 |
| VCA0554 | 3640 | 1975 | 4155 | 1213 | 845 | 1601 | 0,4 | 3,1E-05 | 2,6E-04 |
| VC0731 | 9328 | 18667 | 16354 | 5035 | 7322 | 4748 | 0,4 | 7,7E-05 | 5,8E-04 |
| VC0034 | 1465 | 2168 | 3551 | 936 | 1054 | 1174 | 0,5 | 1,2E-04 | 8,2E-04 |

**Table S1. Genes differentially regulated by indole.**

| | | | | | | | | | |
|---|---|---|---|---|---|---|---|---|---|
| VC0573 | 4031 | 5106 | 3159 | 2130 | 1225 | 2439 | 0,5 | 1,4E-04 | 9,8E-04 |
| VC1440 | 803 | 1073 | 578 | 467 | 268 | 424 | 0,5 | 1,7E-04 | 1,1E-03 |
| VCA0907 | 1430 | 2408 | 1943 | 1139 | 688 | 868 | 0,5 | 2,0E-04 | 1,3E-03 |
| VC2089 | 762 | 1693 | 896 | 567 | 307 | 554 | 0,5 | 2,2E-04 | 1,4E-03 |
| VC1441 | 3956 | 4982 | 2505 | 2010 | 1104 | 2188 | 0,5 | 3,7E-04 | 2,2E-03 |
| VCA0496 | 1389 | 942 | 2071 | 426 | 1108 | 411 | 0,5 | 4,2E-03 | 1,6E-02 |
| Sugar metabolism | | | | | | | | | |
| VCA0896 | 3304 | 3007 | 3497 | 889 | 693 | 872 | 0,3 | 1,6E-32 | 1,8E-30 |
| VC1126 | 7293 | 10250 | 7992 | 2766 | 2625 | 2583 | 0,3 | 2,4E-18 | 1,2E-16 |
| VCA0657 | 685 | 529 | 732 | 221 | 201 | 201 | 0,3 | 7,2E-14 | 2,5E-12 |
| VC1004 | 6034 | 4770 | 6826 | 1934 | 1231 | 1837 | 0,3 | 4,2E-13 | 1,4E-11 |
| VC2024 | 3794 | 3293 | 4602 | 1688 | 1965 | 1679 | 0,5 | 4,2E-09 | 8,0E-08 |
| VCA0897 | 3467 | 2804 | 1622 | 845 | 579 | 943 | 0,3 | 4,9E-09 | 8,9E-08 |
| VCA0898 | 5767 | 4943 | 6369 | 2709 | 1746 | 2587 | 0,4 | 7,2E-09 | 1,3E-07 |
| VC0910 | 3175 | 3395 | 3392 | 1601 | 1326 | 1539 | 0,5 | 1,1E-05 | 1,0E-04 |
| VC0911 | 4479 | 3736 | 4242 | 1881 | 2225 | 1776 | 0,5 | 2,5E-05 | 2,2E-04 |
| VC2480 | 1448 | 1607 | 2584 | 982 | 741 | 885 | 0,5 | 5,1E-05 | 4,0E-04 |
| VCA1060 | 2171 | 771 | 1674 | 633 | 563 | 664 | 0,4 | 1,4E-04 | 9,5E-04 |
| VC2183 | 5155 | 4222 | 6751 | 2904 | 1660 | 3070 | 0,5 | 2,8E-04 | 1,7E-03 |
| Aminoacid metabolism | | | | | | | | | |
| VC1658 | 2116 | 1124 | 2478 | 185 | 194 | 252 | 0,1 | 8,2E-26 | 7,4E-24 |
| VC0008 | 724 | 505 | 822 | 117 | 118 | 140 | 0,2 | 1,2E-20 | 7,9E-19 |
| VC0009 | 679 | 385 | 742 | 88 | 83 | 99 | 0,2 | 4,2E-20 | 2,6E-18 |
| VC0472 | 6447 | 7377 | 9080 | 2458 | 1906 | 2154 | 0,3 | 3,0E-19 | 1,7E-17 |
| VC1312 | 2358 | 2539 | 3364 | 624 | 800 | 669 | 0,3 | 1,9E-18 | 9,8E-17 |
| VC0010 | 3040 | 2440 | 3330 | 550 | 251 | 670 | 0,2 | 2,0E-14 | 7,7E-13 |
| VC0880 | 758 | 657 | 1076 | 241 | 122 | 270 | 0,3 | 2,2E-10 | 5,2E-09 |
| VCA1073 | 509 | 701 | 623 | 250 | 182 | 291 | 0,4 | 3,3E-09 | 6,3E-08 |
| VC0743 | 6022 | 4953 | 6483 | 2669 | 1673 | 2698 | 0,4 | 7,6E-08 | 1,1E-06 |
| VC1995 | 1199 | 1731 | 2309 | 639 | 415 | 724 | 0,4 | 8,9E-08 | 1,3E-06 |
| VCA0815 | 3885 | 3098 | 3980 | 1836 | 1216 | 1630 | 0,4 | 1,7E-07 | 2,4E-06 |
| VC0907 | 596 | 504 | 737 | 298 | 182 | 300 | 0,4 | 2,0E-06 | 2,3E-05 |
| VC0941 | 10076 | 7756 | 8318 | 4801 | 2914 | 4531 | 0,5 | 2,3E-06 | 2,6E-05 |
| VC2682 | 2219 | 1588 | 1853 | 772 | 900 | 1076 | 0,5 | 5,2E-06 | 5,2E-05 |
| VC0947 | 10671 | 13096 | 9872 | 4646 | 6635 | 4962 | 0,5 | 1,5E-05 | 1,4E-04 |
| VCA0772 | 607 | 666 | 968 | 364 | 202 | 431 | 0,5 | 8,6E-05 | 6,4E-04 |
| VC0275 | 3965 | 3080 | 4200 | 1403 | 423 | 1669 | 0,4 | 1,0E-04 | 7,2E-04 |
| VC2485 | 1370 | 3351 | 2132 | 1019 | 971 | 914 | 0,5 | 1,1E-04 | 8,0E-04 |
| VCA0088 | 2895 | 3412 | 2191 | 1471 | 647 | 1654 | 0,5 | 3,7E-04 | 2,2E-03 |
| VC0662 | 1746 | 1104 | 2625 | 973 | 890 | 798 | 0,5 | 3,8E-04 | 2,2E-03 |
| VC0576 | 3097 | 3766 | 9997 | 1913 | 2839 | 1831 | 0,4 | 7,3E-04 | 3,9E-03 |
| VCA0076 | 970 | 1503 | 1188 | 541 | 146 | 612 | 0,4 | 9,4E-04 | 4,8E-03 |

**Table S1. Genes differentially regulated by indole.**

| | | | | | | | | | |
|---|---|---|---|---|---|---|---|---|---|
| VC1492 | 2786 | 6743 | 2369 | 1909 | 1011 | 2088 | 0,5 | 1,5E-03 | 6,9E-03 |
| VC2746 | 19210 | 29045 | 38036 | 10763 | 19348 | 10475 | 0,5 | 2,3E-03 | 9,8E-03 |
| Nucleotide metabolism | | | | | | | | | |
| VC0276 | 5517 | 4824 | 5808 | 1963 | 752 | 2223 | 0,3 | 3,1E-06 | 3,3E-05 |
| VC0767 | 8972 | 9299 | 12691 | 4208 | 4262 | 4502 | 0,4 | 2,4E-10 | 5,7E-09 |
| VC0768 | 17636 | 14237 | 15989 | 5714 | 3303 | 5613 | 0,3 | 1,3E-09 | 2,7E-08 |
| VC0986 | 16980 | 16609 | 14315 | 7288 | 6361 | 7172 | 0,4 | 9,9E-09 | 1,7E-07 |
| VC1228 | 2458 | 2040 | 2902 | 784 | 365 | 922 | 0,3 | 1,6E-07 | 2,2E-06 |
| VC1721 | 2424 | 2857 | 3565 | 1020 | 1480 | 1120 | 0,4 | 3,7E-08 | 6,0E-07 |
| VC2171 | 1886 | 1374 | 2592 | 1049 | 478 | 1191 | 0,5 | 3,2E-03 | 1,3E-02 |
| VC2227 | 7683 | 7434 | 9542 | 1542 | 600 | 1786 | 0,2 | 3,4E-12 | 9,6E-11 |
| VC2258 | 5189 | 4353 | 6703 | 2373 | 3004 | 1852 | 0,5 | 1,3E-06 | 1,6E-05 |
| VC2277 | 6821 | 6135 | 7227 | 1701 | 1908 | 1914 | 0,3 | 1,0E-18 | 5,3E-17 |
| VC2352 | 8009 | 5520 | 8212 | 3958 | 2318 | 4075 | 0,5 | 3,7E-04 | 2,2E-03 |
| VC2712 | 6475 | 3650 | 5923 | 2192 | 772 | 2789 | 0,4 | 7,2E-04 | 3,8E-03 |
| VCA0607 | 1153 | 603 | 1425 | 331 | 773 | 334 | 0,5 | 3,7E-03 | 1,5E-02 |
| VC1365 | 972 | 1192 | 1204 | 492 | 581 | 539 | 0,5 | 5,0E-07 | 6,5E-06 |
| Translation | | | | | | | | | |
| VC0218 | 57141 | 36230 | 62003 | 25376 | 23118 | 26376 | 0,5 | 6,6E-06 | 6,5E-05 |
| VC0291 | 10456 | 11726 | 12645 | 5607 | 4153 | 5227 | 0,4 | 6,1E-11 | 1,5E-09 |
| VC0359 | 62186 | 45942 | 57114 | 27307 | 22476 | 29864 | 0,5 | 1,1E-05 | 1,0E-04 |
| VC0360 | 65435 | 46613 | 78079 | 26811 | 25570 | 27739 | 0,4 | 4,5E-07 | 6,0E-06 |
| VC0361 | 29304 | 38455 | 33303 | 17321 | 16307 | 16634 | 0,5 | 4,4E-10 | 9,8E-09 |
| VC0369 | 24665 | 21200 | 59251 | 15852 | 6855 | 16988 | 0,4 | 1,8E-03 | 8,1E-03 |
| VC0379 | 793 | 970 | 1720 | 567 | 485 | 531 | 0,5 | 4,7E-05 | 3,8E-04 |
| VC0520 | 15942 | 19754 | 24705 | 6763 | 8291 | 6571 | 0,4 | 9,5E-09 | 1,6E-07 |
| VC0570 | 55026 | 72405 | 87623 | 29769 | 23007 | 32471 | 0,4 | 1,1E-08 | 1,9E-07 |
| VC0571 | 20538 | 27307 | 71337 | 12432 | 9502 | 11204 | 0,3 | 2,1E-06 | 2,4E-05 |
| VC0644 | 1515 | 1015 | 1370 | 552 | 434 | 509 | 0,4 | 1,3E-09 | 2,7E-08 |
| VC0645 | 2212 | 1491 | 2617 | 645 | 503 | 659 | 0,3 | 2,2E-12 | 6,5E-11 |
| VC0646 | 74425 | 45508 | 52075 | 24930 | 23318 | 30660 | 0,5 | 1,7E-05 | 1,5E-04 |
| VC0659 | 4553 | 4864 | 6873 | 1701 | 1920 | 1491 | 0,3 | 3,0E-14 | 1,1E-12 |
| VC1209 | 8390 | 9867 | 9418 | 4451 | 4396 | 4138 | 0,5 | 6,6E-11 | 1,7E-09 |
| VC1219 | 1824 | 1726 | 2168 | 991 | 864 | 970 | 0,5 | 1,6E-10 | 3,8E-09 |

**Table S1. Genes differentially regulated by indole.**

| | | | | | | | | | |
|---|---|---|---|---|---|---|---|---|---|
| VC1220 | 4783 | 4380 | 6074 | 2824 | 2175 | 2437 | 0,5 | 1,4E-07 | 1,9E-06 |
| VC1297 | 5980 | 5708 | 9049 | 2891 | 2950 | 3392 | 0,5 | 5,1E-08 | 8,1E-07 |
| VC1640 | 42672 | 27586 | 45907 | 10871 | 10746 | 14654 | 0,3 | 8,3E-09 | 1,5E-07 |
| VC1923 | 14635 | 30088 | 14378 | 9167 | 11082 | 8555 | 0,5 | 7,1E-04 | 3,8E-03 |
| VC2025 | 10223 | 7808 | 14105 | 3020 | 3736 | 2634 | 0,3 | 1,5E-09 | 3,0E-08 |
| VC2026 | 69458 | 52676 | 55871 | 21480 | 25695 | 22013 | 0,4 | 1,2E-07 | 1,7E-06 |
| VC2214 | 6055 | 7838 | 8737 | 3254 | 3998 | 2586 | 0,5 | 1,1E-06 | 1,4E-05 |
| VC2259 | 34754 | 44090 | 37821 | 18907 | 21812 | 17310 | 0,5 | 4,8E-07 | 6,2E-06 |
| VC2342 | 66170 | 86994 | 100616 | 45347 | 39129 | 36577 | 0,5 | 5,5E-08 | 8,5E-07 |
| VC2461 | 2014 | 1761 | 2572 | 814 | 837 | 695 | 0,4 | 9,0E-13 | 2,9E-11 |
| VC2579 | 8782 | 23257 | 18636 | 7681 | 5267 | 7624 | 0,4 | 7,6E-05 | 5,7E-04 |
| VC2583 | 8239 | 20804 | 14710 | 7616 | 6339 | 6607 | 0,5 | 2,7E-04 | 1,7E-03 |
| VC2620 | 1574 | 1292 | 1490 | 785 | 527 | 762 | 0,5 | 2,2E-05 | 1,9E-04 |
| VC2664 | 4009 | 5269 | 4052 | 1503 | 1695 | 1142 | 0,3 | 4,5E-14 | 1,6E-12 |
| Others | | | | | | | | | |
| csrC2 | 3606 | 1920 | 2667 | 508 | 802 | 551 | 0,3 | 9,0E-11 | 2,2E-09 |
| VIBCH10294 | 454 | 271 | 1091 | 284 | 121 | 291 | 0,4 | 5,8E-03 | 2,1E-02 |
| VC0033 | 661 | 587 | 1321 | 282 | 403 | 265 | 0,4 | 1,3E-05 | 1,2E-04 |
| VC1003 | 4970 | 3770 | 3744 | 1838 | 1149 | 2060 | 0,4 | 8,1E-07 | 1,0E-05 |
| VC1490 | 685 | 850 | 1294 | 330 | 344 | 399 | 0,4 | 1,4E-06 | 1,6E-05 |
| VC1208 | 3478 | 4196 | 5582 | 2153 | 1811 | 1986 | 0,5 | 1,4E-07 | 2,0E-06 |
| VC2460 | 3572 | 3075 | 3149 | 1084 | 1300 | 1026 | 0,4 | 2,8E-20 | 1,8E-18 |
| VC0290 | 7761 | 7686 | 8352 | 2994 | 2610 | 3228 | 0,4 | 5,3E-16 | 2,4E-14 |
| VC2459 | 960 | 791 | 658 | 428 | 353 | 392 | 0,5 | 8,0E-08 | 1,2E-06 |
| VC2647 | 6843 | 4552 | 3914 | 1936 | 3109 | 2207 | 0,5 | 1,8E-04 | 1,2E-03 |
| VC0749 | 1248 | 1409 | 1596 | 739 | 601 | 674 | 0,5 | 3,0E-10 | 6,7E-09 |
| VC0766 | 1307 | 1242 | 3902 | 984 | 737 | 1256 | 0,5 | 6,3E-03 | 2,2E-02 |
| Membrane - transporters | | | | | | | | | |
| VC1577 | 6497 | 2204 | 6096 | 403 | 1657 | 363 | 0,2 | 1,4E-06 | 1,7E-05 |
| VC1195 | 2511 | 3007 | 2599 | 1076 | 1138 | 1086 | 0,4 | 4,1E-14 | 1,5E-12 |
| VC1962 | 1002 | 693 | 894 | 237 | 323 | 230 | 0,3 | 3,0E-11 | 7,8E-10 |
| VC1043 | 20149 | 34658 | 22588 | 7960 | 8029 | 9141 | 0,3 | 5,1E-09 | 9,3E-08 |
| VCA0862 | 718 | 539 | 837 | 239 | 239 | 338 | 0,4 | 4,9E-08 | 7,8E-07 |
| VC1655 | 1156 | 809 | 1960 | 372 | 425 | 333 | 0,3 | 6,9E-09 | 1,2E-07 |
| VC1409 | 885 | 311 | 759 | 79 | 245 | 73 | 0,3 | 1,1E-06 | 1,3E-05 |
| VC1329 | 2576 | 1202 | 2399 | 450 | 273 | 641 | 0,3 | 5,4E-09 | 9,9E-08 |
| VC2305 | 5931 | 3437 | 9844 | 1711 | 2267 | 2010 | 0,3 | 1,1E-06 | 1,4E-05 |

**Table S1. Genes differentially regulated by indole.**

| | | | | | | | | | |
|---|---|---|---|---|---|---|---|---|---|
| VCA0554 | 3640 | 1975 | 4155 | 1213 | 845 | 1601 | 0,4 | 3,1E-05 | 2,6E-04 |
| VC2278 | 7821 | 6452 | 9648 | 2026 | 1781 | 2313 | 0,3 | 3,4E-18 | 1,7E-16 |
| VC1319 | 3625 | 988 | 2722 | 474 | 1989 | 485 | 0,5 | 1,6E-02 | 4,9E-02 |
| VCA1071 | 777 | 918 | 739 | 333 | 197 | 313 | 0,4 | 1,5E-11 | 4,0E-10 |
| VC1669 | 764 | 292 | 1161 | 137 | 223 | 135 | 0,3 | 1,4E-07 | 1,9E-06 |
| VC0156 | 13036 | 12336 | 11471 | 2758 | 7063 | 2800 | 0,4 | 6,7E-05 | 5,1E-04 |
| VC0475 | 1465 | 834 | 1238 | 579 | 468 | 527 | 0,5 | 1,5E-06 | 1,8E-05 |
| Hyp | | | | | | | | | |
| VC0268 | 1534 | 1816 | 2463 | 739 | 1253 | 701 | 0,5 | 5,3E-04 | 3,0E-03 |
| VC0519 | 67392 | 74591 | 56109 | 23575 | 31849 | 20968 | 0,4 | 1,0E-07 | 1,5E-06 |
| VC0714 | 3388 | 1594 | 2591 | 663 | 2020 | 811 | 0,5 | 8,5E-03 | 2,9E-02 |
| VC1052 | 1431 | 1646 | 2032 | 291 | 395 | 315 | 0,2 | 7,1E-20 | 4,3E-18 |
| VC1058 | 762 | 936 | 1507 | 368 | 638 | 343 | 0,5 | 1,0E-03 | 5,2E-03 |
| VC1074 | 639 | 609 | 1692 | 286 | 317 | 303 | 0,3 | 2,5E-06 | 2,8E-05 |
| VC1317 | 4580 | 788 | 3907 | 213 | 1306 | 301 | 0,4 | 6,8E-03 | 2,4E-02 |
| VC1574 | 1767 | 647 | 1134 | 366 | 505 | 432 | 0,4 | 7,6E-06 | 7,3E-05 |
| VC1575 | 786 | 269 | 1065 | 134 | 207 | 119 | 0,3 | 1,0E-07 | 1,5E-06 |
| VC1576 | 2727 | 1008 | 1653 | 330 | 786 | 345 | 0,3 | 2,5E-06 | 2,8E-05 |
| VC1578 | 3800 | 1491 | 1767 | 278 | 1236 | 270 | 0,3 | 2,3E-04 | 1,5E-03 |
| VC1832 | 1010 | 861 | 1883 | 390 | 605 | 473 | 0,4 | 4,6E-05 | 3,7E-04 |
| VC1941 | 3896 | 3191 | 4799 | 1591 | 1861 | 1619 | 0,4 | 1,5E-08 | 2,5E-07 |
| VC2093 | 364 | 468 | 980 | 286 | 316 | 262 | 0,5 | 1,2E-03 | 6,0E-03 |
| VC2443 | 1378 | 1147 | 2022 | 611 | 768 | 809 | 0,5 | 4,3E-05 | 3,5E-04 |
| VC2472 | 1677 | 1930 | 2799 | 671 | 813 | 648 | 0,4 | 9,3E-10 | 1,9E-08 |
| VC2706 | 14791 | 12371 | 19055 | 2652 | 2846 | 3038 | 0,2 | 2,8E-28 | 2,8E-26 |
| VCA0026 | 2034 | 2585 | 2667 | 1005 | 1465 | 984 | 0,5 | 7,5E-05 | 5,6E-04 |
| VCA0921 | 4725 | 4659 | 8190 | 1557 | 2134 | 1754 | 0,3 | 8,6E-09 | 1,5E-07 |
| Metabol | | | | | | | | | |
| VC0522 | 4113 | 2988 | 3708 | 1695 | 2094 | 1487 | 0,5 | 8,2E-07 | 1,0E-05 |
| VC1942 | 1302 | 1679 | 3645 | 440 | 438 | 455 | 0,2 | 2,0E-11 | 5,3E-10 |
| VC0869 | 14734 | 12746 | 18061 | 3007 | 1208 | 3242 | 0,2 | 2,7E-12 | 7,7E-11 |
| VC1040 | 977 | 1202 | 1356 | 697 | 434 | 627 | 0,5 | 8,6E-06 | 8,3E-05 |
| VCA0150 | 2635 | 1976 | 2642 | 1116 | 702 | 1170 | 0,4 | 1,4E-05 | 1,3E-04 |
| VCA0614 | 2154 | 1520 | 2118 | 759 | 457 | 842 | 0,4 | 1,2E-07 | 1,7E-06 |
| VCA0558 | 2076 | 607 | 4525 | 607 | 1460 | 718 | 0,5 | 1,0E-02 | 3,4E-02 |
| VCA0496 | 1389 | 942 | 2071 | 426 | 1108 | 411 | 0,5 | 4,2E-03 | 1,6E-02 |
| VC0992 | 1408 | 1336 | 1672 | 627 | 435 | 786 | 0,4 | 1,1E-07 | 1,6E-06 |
| VCA0843 | 1854 | 2880 | 1583 | 839 | 200 | 997 | 0,4 | 4,7E-04 | 2,7E-03 |
| VC2545 | 10175 | 14016 | 15506 | 3639 | 4482 | 3767 | 0,3 | 1,8E-10 | 4,4E-09 |
| VC0745 | 15244 | 15708 | 17276 | 6155 | 7574 | 7144 | 0,4 | 3,3E-08 | 5,3E-07 |

**Table S1. Genes differentially regulated by indole.**

| | | | | | | | | | |
|---|---|---|---|---|---|---|---|---|---|
| VCA0863 | 2047 | 1494 | 2025 | 616 | 561 | 814 | 0,4 | 3,9E-11 | 1,0E-09 |
| VCA0661 | 794 | 886 | 1062 | 483 | 357 | 513 | 0,5 | 2,9E-06 | 3,1E-05 |
| VC0051 | 4018 | 2862 | 4106 | 1016 | 374 | 1138 | 0,3 | 5,6E-08 | 8,7E-07 |
| VC0052 | 2282 | 2082 | 3109 | 621 | 263 | 597 | 0,2 | 8,0E-12 | 2,2E-10 |
| VC2226 | 11858 | 12602 | 16080 | 2538 | 1208 | 2848 | 0,2 | 3,7E-15 | 1,5E-13 |
| VC1625 | 6073 | 5424 | 5715 | 1808 | 1036 | 2011 | 0,3 | 1,3E-12 | 3,8E-11 |
| VC1738 | 8039 | 9711 | 16080 | 6283 | 5005 | 4652 | 0,5 | 4,8E-05 | 3,8E-04 |
| VC2256 | 2868 | 2130 | 4467 | 1588 | 1450 | 1314 | 0,5 | 2,5E-05 | 2,1E-04 |

Table S2. Strains, plasmids and primers

| **Strain** | **Strain number** | **Construction** |
|---|---|---|
| ***Vibrio cholerae*** | | |
| N16961 wt strain | | |
| N16961 hapR+ wt strain | F606 | Gift from Melanie Blokesch |
| *ΔraiA (VC0706)* | J251 | PCR amplification of 500bp up and down regions of VC0706 using primers ZIP381/384 and ZIP382/383. PACR amplicifaction of aadA7 conferring spectinomycin resistance on pAM34 using ZB47/48. PCR assembly of the VCA0706::spec fragment using ZIP383/384 and allelic exchange by natural transformation, , as described previously (Val et al PLoS Genetics 2012, Negro et al, mBio 2019) |
| *Δrmf (VC1484)* | L557 | allelic exchange by integration and excision of conjugative suicide plasmid pMP7 L045, replacing the gene with *frt::kan::frt* as described previously (Val et al PLoS Genetics 2012, Negro et al, mBio 2019) |
| *Δhpf (VC2530)* | M566 | allelic exchange by integration and excision of conjugative suicide plasmid pMP7 M472, replacing the gene with frt::kan::frt as described previously (Val et al PLoS Genetics 2012, Negro et al, mBio 2019) |
| *ΔraiA Δrmf* | L790 | allelic exchange by integration and excision of conjugative suicide plasmid pMP7 L045 in J251, replacing the gene with frt::kan::frt as described previously (Val et al PLoS Genetics 2012, Negro et al, mBio 2019) |
| *Δcrp* | 9950 | allelic deletion by integration and excision of conjugative suicide plasmid pMP7 8348 , replacing the gene with spec |
| *Δfur* | N541 | allelic exchange by integration and excision of conjugative suicide plasmid pMP7 N450, replacing the gene with frt::kan::frt as described previously (Val et al PLoS Genetics 2012, Negro et al, mBio 2019) |
| *ΔrpoS* | A321 | *Baharoglu et al, PLoS Genetics 2013* |
| *ΔtnaA (VC0161)* | J253 | PCR amplification of 500bp up and down regions of VC0161 using primers ZIP93/94 and ZIP95/96. PACR amplicifaction of aadA7 conferring spectinomycin resistance on pAM34 using ZIP97/98. PCR assembly of the VCA0161::spec fragment using ZIP93/96 and allelic exchange by natural transformation. |
| ***Escherichia coli*** | | |
| MG1655 wt strain | | laboratory collection |
| *ΔraiA* | J237 | P1 transduction from KEIO strain JW2578-1 |
| ***Pseudomonas aeruginosa* PAO1** | | laboratory collection |
| **Plasmids** | | |
| pBAD43 | | spectinomycin resistant. Carries Para promoter repressed by glucose 1% and induced by arabinose 0.2%. (Guzman, 1995) |
| pBAD43-RaiA+ | L907 | *raiA* (VC0706) amplified using primers raiAeco/raiAxba and cloned under Pbad using 5' EcoRI and 3' XbaI restriction sites |
| pBAD43-Hpf+ | M834 | *hpf* (VC2530) amplified using primers hpfeco/hpfxba and cloned under Pbad promoter using 5' EcoRI and 3' XbaI restriction sites |

| | | |
|---|---|---|
| pBAD43-Rmf+ | M831 | *rmf* (VC1484) amplified using primers rmfeco/rmfxba and cloned under Pbad using 5' EcoRI and 3' XbaI restriction sites |
| pSC101-PraiA-gfp | L707 | raiA promoter region (500bp) was amplified using primers ZIP383/446 and gfp was amplified using primers zip443/444. 2 fragments were assembled by PCR assembly using primers ZIP383/ZIP443, so that *gfp* is fused to the *raiA* promoter at the ATG start codon of *raiA*. The assembled fragment was cloned into pTOPO and extracted using EcoRI and cloned into pSC101 low copy plasmid (carbenicillin resistant). |
| pMP7-Δhpf::kan | M472 | gibson assembly using primers MV450/451 for the amplification of pMP7 vector, primers VC2530hpf5bis/7 and VC2530hpf6bis/8bis for up and down regions of the gene, and primers MV268/269 on pKD4 plasmid for the resistance gene (frt::kan::frt). |
| pMP7-Δrmf::kan | L045 | gibson assembly using primers MV450/451 for the amplification of pMP7 vector, primers VC1484rmf5/7 and VC1484rmf6/8 for up and down regions of the gene, and primers MV268/269 on pKD4 plasmid for the resistance gene (frt::kan::frt). |
| pMP7-Δcrp | 8348 | (ref: Baharoglu et al, J bact 2012) |
| pMP7-Δfur::kan | N450 | gibson assembly using primers MV450/451 for the amplification of pMP7 vector, primers VC2106fur5/7 and VC2106fur9/8 for up and down regions of the gene, and primers MV268/269 on pKD4 plasmid for the resistance gene (frt::kan::frt). |
| **primers** | | |
| raiAecoRI | | ggaattcaccATGAAAATCAACATCACTGGTAA |
| raiAxba | | gctctagaTTATTCCACTTCTTCGCTCAG |
| hpfeco | | ggaattcaccATGCAAATCAACATTCAAGGCC |
| hpfxba | | gctctagaTTAATGACTACTTAGCTTTTCTTT |
| rmfeco | | ggaattcaccATGAAGAGACAAAAGCGTGAT |
| rmfxba | | gctctagaTTATTTGCAGAGACCAGAAAGTTT |
| ZB47 | | CCCGTTCCATACAGAAGCTGGGCGAACAAACGATGCTCGC |
| ZB48 | | GACATTATTTGCCGACTACCTTGGTGATCTCGCCTTTCACG |
| ZIP381 | | GCGAGCATCGTTTGTTCGCCCAGCTTCTGTATGGAACGGGTAAATAGAATG ATGGGAGATAGCGC |
| ZIP382 | | CGTGAAAGGCGAGATCACCAAGGTAGTCGGCAAATAATGTCAATGCTTTTT CCTCTGTGTCATCCCTTATGG |
| ZIP383 | | AACCTATCCAGATACCGAAGCGGC |
| ZIP384 | | GTTTTGCAGATAGGATTGCTGAAGC |
| ZIP93 | | GTCGAATGCCAAACGAAAGCGGAAAATACC |
| ZIP94 | | GCGAGCATCGTTTGTTCGCCCAGCTTCTGTATGGAACGGGTACGGTATTGA AAAAGTGCAGCCTGC |
| ZIP95 | | CGTGAAAGGCGAGATCACCAAGGTAGTCGGCAAATAATGTCGCTGGGTAT TCCTAAAAATTAAAATATAAAGTCATGC |
| ZIP96 | | CACAACAACTCTAGTATCTGGTTTACCCTCG |
| ZIP97 | | GGCAGCCTATGCAGGCTGCACTTTTTCAATACCGTCCCGTTCCATACAGAA GCTGGGCGAACAAACGATGCTCGC |
| ZIP98 | | ATGACTTTATATTTTAATTTTTAGGAATACCCAGCGACATTATTTGCCGACTA CCTTGGTGATCTCGCCTTTCACG |
| ZIP443 | | TTATTTGTATAGTTCATCCATGCCATGTGTAATCCCAGC |
| ZIP444 | | ATGCGTAAAGGAGAAGAACTTTTCACTGGAGTTGTCC |

| | | |
|---|---|---|
| ZIP446 | | GGACAACTCCAGTGAAAAGTTCTTCTCCTTTACGCATAATGCTTTTTCCTCT GTGTCATCCCTTATGG |
| VC2530hpf5bis | | CTATTATTTAAACTCTTTCCgccgatggtgctcaacgatg |
| VC2530hpf7 | | CTACACAATCGCTCAAGACGTGagactttccttctctagtttagg |
| VC2530hpf6bis | | TACGTAGAATGTATCAGACTcttcacactgcaatagcacg |
| VC2530hpf8bis | | CTAATTCCCATGTCAGCCGTGCGCCAGATCGCCGAAGCTA |
| VC1484rmf5 | | CTATTATTTAAACTCTTTCCggcttggttgttagatgatc |
| VC1484rmf7 | | CTACACAATCGCTCAAGACGTGagttctgtcctcatacggta |
| VC1484rmf6 | | TACGTAGAATGTATCAGACTgttgatgatggatcgcattacc |
| VC1484rmf8 | | CTAATTCCCATGTCAGCCGTttcccatacaacaaagcttga |
| VC2106fur5 | | CTATTATTTAAACTCTTTCCAAGCGGATGCGAACTTCGC |
| VC2106fur7 | | CTACACAATCGCTCAAGACGTGATACTTTCCTGTTGATGTTCTGC |
| VC2106fur8 | | CTAATTCCCATGTCAGCCGTGCTCACAAGCCGAAGAAATAA |
| VC2106fur9 | | TACGTAGAATGTATCAGACTccacaaatcgatcagtttatgg |
